# Electrically programmable picoscale phototransduction of a newly discovered microbial rhodopsin

**DOI:** 10.64898/2026.05.29.728716

**Authors:** Ilaria Cardace, Lorenzo Dominici, Vincenzo Ardizzone, Adriano Cola, Antonio Fieramosca, Concetta Nobile, Fabio Polticelli, Fabio Bruni, Milena De Giorgi, Dario Ballarini, Giuseppe Gigli, Luisa De Marco, Daniele Sanvitto

## Abstract

Human retina can achieve single-photon sensitivity through specialised photoreceptors that convert light into electrical signals via phototransduction. Among microbial light-sensitive proteins, proteorhodopsins stand out for their intrinsic light-driven ion transport and spectral tunability, making them promising candidates for bio-inspired photonic devices. A central challenge for acellular integration, however, is the fragility of most bacterial rhodopsins under extreme conditions. Here, we exploit the exceptional robustness of TARA76, a microbial rhodopsin that retains structural integrity even upon complete dehydration, to demonstrate its functional reconstitution in an artificial black lipid membrane within a biocompatible microfluidic platform. By recording light-induced ionic currents with picoampere sensitivity across a broad range of pH, illumination power, electrolyte composition, and applied voltages, we establish TARA76 as a high-performance photoelectric transducer in a fully acellular environment. Strikingly, we uncover a strong and previously unreported dependence of the photocurrent on Na^+^ ions, which appears to play a key structural and functional role in stabilising the protein’s active conformation. Furthermore, we demonstrate that the orientation of TARA76 within the artificial membrane can be externally controlled by applying a defined electric field during bilayer formation, enabling deterministic tuning of photocurrent directionality. Together, these results establish a robust and miniaturisable bio-photonic platform with direct implications for quantum light sensing, neuromorphic bioelectronics, and next-generation artificial retinal interfaces.

## Introduction

Genetically encoded, light-responsive proteins can control biological events at the level of single cells or entire organisms. In nature, specialised cells can absorb a single photon with remarkable efficiency.^1,2^ For instance, the human retina employs photoreceptor cells to mediate vision through a phototransduction process that converts light energy into electrical signals. These signals are regulated by various stimuli, such as changes in voltage, light, and ligand binding, which can activate specific ion channels in the plasma membrane, thereby triggering intracellular enzymatic pathways.^1–3^

In this field, specific photoreceptors undergo light-induced conformational changes that enable the selective movement of ions across the plasma membrane.^4–6^ Based on their biological origin, these proteins are grouped into type I (microbial) and type II (animal) rhodopsins. They are essential for environmental adaptation,^7–9^ and can differ in their mechanism of action, such as outward proton pumps^10^, inward proton pumps^11^ and cation channels.^5,12,13^ Understanding the functionality and sensitivity of these proteins is crucial, as they have applications in various fields, ranging from biology to next-generation optoelectronics and hybrid devices. One of the fields in which microbial rhodopsins are most used is optogenetics, where natural opsins are engineered and introduced into non-photosensitive cells.^4,14–16^ When integrated into the neuronal membrane, these opsins modulate the membrane potential in response to light, enabling non-invasive stimulation of action potentials. This technology is revolutionising the study of complex neuronal circuits.^9,17–21^

Beyond optogenetics, a less-exploited but equally interesting field is the use of microbial rhodopsins as highly sensitive light detectors, potentially serving as bio-inspired components in optical neural networks and quantum technologies. Their intrinsic ability to detect single photons could, in fact, be leveraged to build high-performance detectors,^2^ while their nonlinear response could be advantageous for neuromorphic applications. Integrating biological components into electronic devices opens new ways to access, analyse, and respond to intercellular information through data transduction and signal transmission.^22–24^ Most research on biosensors and bioelectronic devices has traditionally focused on bacteriorhodopsin (BR) from *Halobacterium salinarum*,^25^ which serves as a prototypical system and a key benchmark for structural and functional performance. In contrast, proteorhodopsins (PRs) are a more recently discovered, highly diverse class of microbial rhodopsins.^7,26,27^ Due to their exceptional sequence variability and spectral tunability,^7^ these bacterial light-driven outward proton pumps constitute the largest family in the group and, although still minimally investigated for bioelectronic applications, are increasingly recognized as highly promising candidates for next-generation devices.

In this work, we report the successful incorporation of the exceptionally resistant yet largely unexplored TARA76 proteorhodopsin into an artificial lipid membrane and its comprehensive characterisation. The dual objectives pursued were firstly to isolate the TARA76 within an artificial lipid membrane to investigate its intrinsic photophysical behaviour in a fully controlled, acellular environment, and secondly to engineer an ultra-efficient light micro-detector based on biological photoreceptors integrated into an optoelectronic device. We focused our attention on the recently discovered TARA76 protein since one of its most notable features is its remarkable stability in dry conditions; upon dehydration, the chromophore turns bright violet, and upon rehydration, it reverts to its original colour.^28^ This suggests that TARA76 may remain functionally active in dry environments, a feature shared with BR from *Halobacterium halobium*^*29*^, indicating potential applications in semiconductor devices and electrobiological systems.

By eliminating the cellular architecture while preserving phototransduction functionality, we demonstrate that this stable proteorhodopsin can be effectively harnessed within compact artificial devices, paving the way for applications such as enhanced-function artificial retinas. Simultaneously, the acellular configuration enables the exploration of previously unknown photo-electrical properties of TARA76, which until now has been described solely by evaluating its proton-pumping activity in bacterial suspension.^28^ Notably, we reveal that its photocurrent can be modulated by different ionic conditions, with a pronounced dependence on Na^+^ ions, suggesting a broader functional role and opening new avenues for bioelectronic and biophysical investigation. In addition, we demonstrate that a practical strategy to control the directionality of the photocurrent can be achieved by applying a defined electric field during bilayer formation, representing a crucial step toward the engineering of bioelectronic interfaces with well-defined polarity.

## Results and discussion

### Incorporation of Tara76 into an artificial membrane device

TARA76 was selected with precision, and its sequence alignment confirmed the high similarity with other photoactive protein pumps within the proteorhodopsin group (see Supplementary Fig. S1). In accordance with the established approach^28^, TARA76 was expressed, extracted from bacterial membranes and purified using immobilised metal affinity chromatography (IMAC). Biochemical and spectroscopic evidence demonstrated its expression at a high yield and the incorporation of the all-trans retinal (see Supplementary Fig. S2).

To preserve TARA76’s natural photocycle and facilitate its integration into an artificial membrane known as Black Lipid Membrane (BLM) ^30^, the protein was reconstituted in nanodiscs (NDs). These nanometric structures are formed by membrane scaffold proteins (MSPs), which can be characterised by different sizes and combined with several phospholipids (Fig. 1a).^31–33^ Spectroscopic analysis of the reconstituted protein revealed two distinct absorbance peaks: one representing the protein backbone, centred around 280 nm (also visible in the empty nanodiscs, Fig. 1b black dashed line), and the other representing the retinal intercalated on the active site of the protein, centred around 490 nm at pH 6 (Fig. 1b, red solid line). Transmission electron microscopy (TEM) characterization of the isolated TARA76-nanodiscs highlights both a disc-shaped morphology and a mean size of ∼ 9÷12 nm of the particles (Fig. 1c), in accordance with dynamic light scattering (DLS) analysis, thus suggesting a stable TARA76-NDs formation (Supplementary Fig. S2b). Finally, SDS-PAGE gel electrophoresis analysis of samples before and after NDs purification further confirmed successful protein incorporation. The final purified sample contained two distinct bands at ∼25 KDa (TARA76) and ∼29 KDa (MSP1E3D1) only if TARA76 is present in the reaction (Supplementary Fig. S2c).

**Figure 1.**
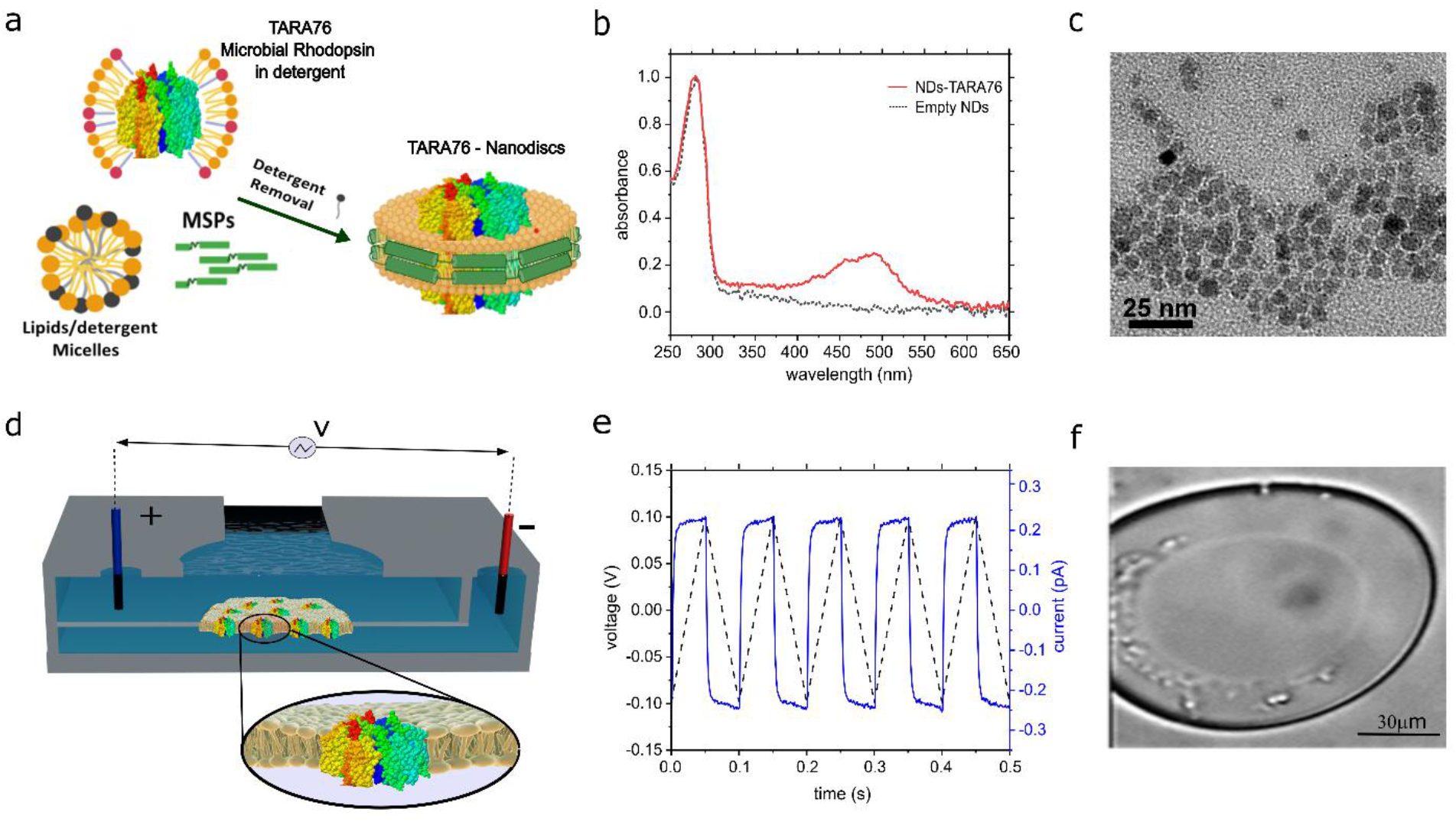
Artificial membrane device. (**a**) Schematic representation of the TARA76 reconstitution into nanodiscs. The reaction mixture consists of the extracted membrane protein TARA76, DMPC lipids in detergent solution, and Membrane Scaffold Proteins (MSP), which provide structural support for the nanodiscs, combined in a defined ratio depending on the size of the protein. Upon detergent removal, spontaneous self-assembly occurs, leading to the formation of the nanodisc structure. (**b**) Functional absorption spectrum of TARA76 reconstituted in nanodiscs in a reaction characterised by 1:126.5:0.5 ratio respectively for MSP: DMPC: TARA76-monomer (TARA76-NDs, red line) and its relative negative control (empty NDs black line). The spectra were taken in the dark at pH 6. (**c**) Representative TEM image of the ND samples (mean size ∼ 9÷12 nm). (**d**) Schematic representation of the microfluidic device. The artificial membrane is formed across a 120 μm hole aperture on a Teflon septum separating the two microchambers, independently accessible and filled with 0.5 M KCl solution. The AgCl electrodes in each chamber enable electrical measurements. White light from the top of the optical microscope is used to visualise the membrane during formation. (**e**) Typical current response of an artificial lipid membrane upon application of a triangular voltage wave (under dark). (**f**) Bright-field image of bilayer membrane formed using the air bubble technique with DPhPC in n-octane (10 mg/ml) in the microfluidic device.

After their characterization the nanodiscs were added to the microfluidic device where the synthetic lipid membrane is produced. We used a model membrane system composed of a single lipid bilayer, which has long served as a foundational platform for studying membrane biophysics.^34–38^ To form the BLM, the microchambers were filled with a 10 mM NaCl solution, buffered or unbuffered, depending on the experiment (Fig. 1d). A small droplet of a phospholipid dissolved in n-octane was micropipetted near the micro-aperture. A spontaneous energy-minimising thinning of the lipidic phase occurs at the interface between the two aqueous phases, ultimately yielding a stable bilayer membrane. The entire microfluidic device is housed within a Faraday cage, and the immersed AgCl electrodes are connected directly to the current amplifier. The membrane formation was first monitored electrically by applying an AC triangular voltage waveform (100 mV amplitude, at 10 Hz frequency, Fig. 1e black dashed line) across the chambers and measuring the resulting current. In the absence of a BLM, the current signal mirrors the triangular input voltage, indicating a pure resistive pathway. Upon successful BLM formation, the system behaves capacitively, and the current response shifts to a rounded square wave (Fig. 1e, blue solid line), confirming the presence of the non-conductive bilayer. An additional check was achieved optically, as the characteristic formation of an annulus (Plateau-Gibbs border) can be observed under an optical inverted microscope (Fig. 1f).

The capacitance *C* was monitored in real time from the amplitude I_m_ of the measured square-wave current response, according to the relationship *C* =*I*_m_/(*dV*/*dt*), where d*V*/d*t* represents the temporal slope of the applied triangular voltage waveform. As the lipid phase progressively thins, a corresponding increase in capacitance is observed, in agreement with *C* = *ϵ* \**A*/*d*, where A is the membrane area, *d* the thickness of the insulating layer, and *ϵ* the dielectric permittivity. The maximum measured capacitance (∼50 pF) is consistent with the expected value for a single lipid bilayer spanning a 120 µm diameter aperture (Ionovation device, see Methods), assuming a relative dielectric constant of 2.2 and a bilayer thickness of 4-5 nm.^39,40^ The associated specific capacitance (∼0.43 µF cm^−2^) agrees well with canonical values for single lipid bilayers.

To benchmark the BLM formation and assess the sensitivity of the electrical measurement setup, Gramicidin A^41^, a 16-amino acid peptide, was used, which in a phospholipid bilayer dimerises, forming an ion pore.^42^ It was added to the upper and lower chambers of the microfluidic device. Current recordings at a constant voltage of 100 mV exhibited stochastic stepwise current transitions of approximately 2 pA, consistent with the opening and closing of individual ion channels (Supplementary Fig. S3).

### Photocurrent response of TARA76

TARA76 electrical behaviour was analysed upon its incorporation into the artificial membrane to determine its photocycle characteristics. Accordingly, a laser beam, aligned via an optical table, is passed through an inverted optical microscope and entered through the objective lens to irradiate the membrane. The photocurrent was measured under a continuous voltage of 100 mV, i.e. in DC configuration, before and after the addition of the protein (Fig. 2a).

**Figure 2.**
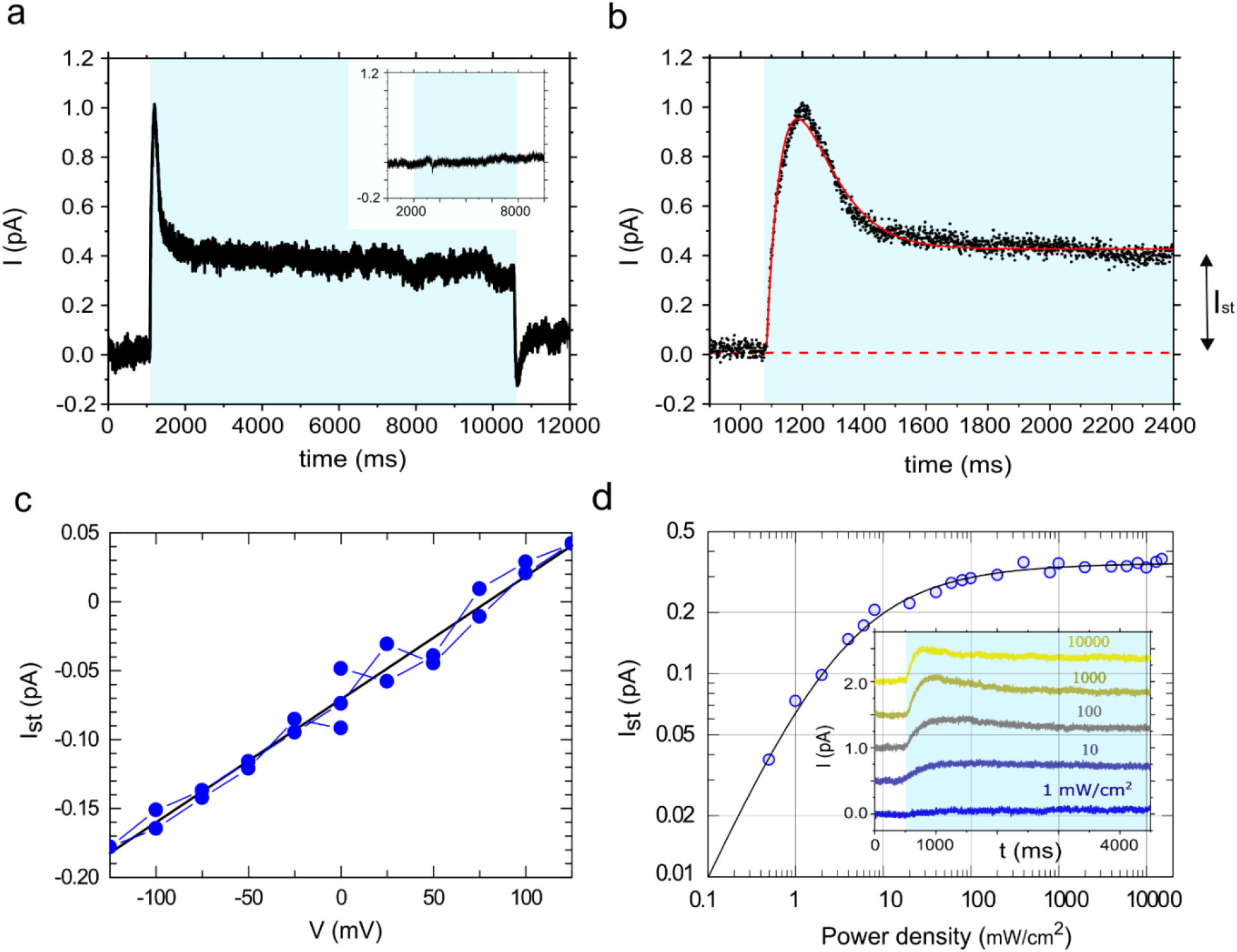
Photocurrent transients. **(a)** Representative current trace from the BLM after the incorporation of TARA76 expressed in *Escherichia coli* (BL21) and reconstituted in nanodiscs. The Illumination time window with a 488 nm laser onto the BLM is indicated by the light blue shaded areas. Inset: current trace before adding TARA76-loaded NDs, showing no response to light. (**b**) Bi-exponential fit (red line) of the current transition (black dots) after the laser switch-on. The I_st_ indicates the amplitude of the long-time stationary current. (**c**) TARA76 light-induced stationary current versus the applied voltage (when scanning V in a closed loop ranging from -125 mV to 125 mV in steps of 25 mV). (**d**) Dependence of the light-induced stationary current versus the laser power density *P* (ranging from a minimum of 0.5 mW cm^-2^ to a maximum of 15 W cm^-2^). Blue dots are experimental data and solid line a guide for the eye. Inset: photocurrent recorded at selected power density (0.001, 0.01, 0.1, 1, 10 W cm^-2^). Experiments in (a,b) and (d) were performed at pH 6 in phosphate-sodium buffer with a constant applied voltage of 100 mV. All the experiments shown here were performed at the concentration of TARA76 equal to 4.3 μM and with a sampling time of 1 ms.

When the laser is switched on in the presence of TARA76, a rapid increase in current is observed, as shown in Fig. 2a (shaded cyan area shows the light exposure time range), indicating the activation of a light-gated ion transport mechanism. A successive decrease follows the initial current spike, eventually reaching a steady-state level of photocurrent, in agreement with previous studies for different proteorhodopsins.^26,43^ Upon termination of the light stimulus, the current returns to baseline, suggesting that the protein relaxes back to its original, dark-state conformation. As shown in the inset of Fig. 2a, no light-dependent current was observed in the absence of TARA76, confirming that the response is protein dependent. The photocurrent response after laser light activation can be described by one time constant for the rapid increase, τ_1_ = 80 ms, and one time constant for the slightly slower decay, τ_2_ = 85 ms (see fit in Fig. 2b). These values are not related to the internal response mechanisms of the membrane but to the limited bandwidth of the setup in the present configuration (about 10 Hz), where a high amplification factor is set to detect sub-pA current levels. The initial transient is attributed to rapid electrogenic charge movements, primarily due to retinal isomerisation and early proton transfer events within the proteorhodopsins photocycle.^44^ It precedes the establishment of steady-state proton pumping and serves as a hallmark of the dynamic conformational changes that occur in the light-activated state of the protein.^11,45–47^ The residual level of the current at long time (a few seconds) is labelled as the stationary current I_st_. The I_st_(λ) response was verified versus the irradiation wavelength (Supplementary Fig. S4), showing a good agreement with the absorption tail of the protein (in the accessible spectroscopic range of our setup).

We initially performed experiments varying TARA76 concentration, and the results suggest a saturation behaviour at a value of 4.3 µM (Supplementary Fig. S5). This concentration was then used for all the following measurements. The Fig. 2c shows the behaviour of the I_st_ as a function of the applied voltage at pH 6 (in a voltage closed ramp from 0 V to -125 mV, then to +125 mV and back to 0 V in step of 25 mV). This linear dependence indicates that the TARA76 protein can reverse the direction of the proton flow in response to light, depending on the polarity of the applied electric field.^43,48^ At pH 6, the current– voltage relation appears practically linear, and the intercept with the horizontal axis yields a so-called reversal potential of approximately +75 mV. While such a reversal potential is not fixed, depending on the external conditions, we confirmed that the observed linear slope remains essentially constant across different experiments. It is important to note that this slope is associated with the photoconductance ΔG, as it represents the I_st_/V_DC_ ratio, where I_st_ is the step change in current measured a few seconds after light exposure relative to the dark condition immediately before switch-on. In our system, the ΔG is approximately 1.0 pS.

The power dependence revealed a saturation behaviour as well of the I_st_ with respect to the laser power density *P*, as shown in Fig. 2d. While at low irradiation levels the I_st_(*P*) curve is almost linear, it then becomes highly sublinear, reaching a plateau at 1 W cm^-2^. At higher power density (> 10-50 W cm^-2^), one order of magnitude beyond photoresponse saturation the membrane becomes growingly unstable under laser exposition. From these results, we determined the maximum laser power at which the membrane system reaches full activation (<10 W cm^-2^), enabling standardised conditions for subsequent experiments and allowing us to determine the maximum number of proteins present in the membrane. It is interesting to note that a more complex behaviour emerges when considering the full shape of the transients (see the inset to Fig. 2d). A current peak is absent in the lower-intensity traces, where only a monotonic time transient up to I_st_ is visible. As the laser power increases, the peak in the current becomes detectable with maximum visibility at approximately 1 W cm^-2^, followed by the decaying transient to I_st_. Furthermore, after prolonged illumination or under varying conditions, the initial current peak becomes less pronounced, in agreement with Lörinczi et al.^49^ This suggests a more complex interplay among multiple underlying mechanisms.

To evaluate the membrane’s protein occupancy and its electrical activity, we considered the following. From the literature, we found that a single bacteriorhodopsin pumps one proton with a cycle time of τ_cycle_= 53.7 ms, under saturating light conditions, corresponding to a continuous pumping rate of about 18.6 protons per second.^50^ The electrical current generated by a single protein, due to proton charge transport, is therefore 3 aA. In our experiments, the maximum peak observed in the photocurrent transient is about 1 pA. If the measured current arises entirely from proton pumping by functional proteorhodopsin molecules, the total number of simultaneously active molecules during the transient peak is estimated at approximately 3.35 x 10^5^. To evaluate their density inside the artificial membrane system, we need to consider a typical membrane diameter of 80 μm (Elements device, see Methods), which corresponds to a membrane area of 5027 μm^2^. A cross-check of the active area is possible by using this value together with the specific capacitance retrieved above (∼0.43 µF cm^−2^).^39^ The expected resulting capacitance is 22 pF, in good agreement with what was observed in the relative AC experiments (see the following). The order of magnitude for the density of the active proteins in the membrane is therefore 65 μm^-2^, which is consistent with the experimentally observed range in AFM-based studies.^51^ These values remain reasonably compatible, given the experimental variability in these delicate membrane systems, such as the variable annulus, the presence of different monomers, and possible differences in protein orientation and functional efficiency within the artificial membrane.^51^ Besides, the NDs are known to incorporate a single protein,^33,52–54^ and have a defined diameter of about 12 nm. Therefore, the theoretical maximum number of NDs in the membrane is ∼40 x 10^6^ units and their maximum density ∼8 × 10^3^ µm^-2^ (considering the in-plane densest circle packing factor of 0.9). Consequently, the proportion of the pre-existing membrane replaced by NDs appears to be on the order of 0.8%.

### Behaviour with different pH and electrolytes conditions

After establishing the optimal TARA76 concentration and light power density—based on the highest measured yet stable photocurrents—we characterised its response under varying environmental conditions. One objective was to determine the influence of both the pH and the electrolyte solutions on the TARA76 photocurrent. This is even more significant, since such a protein has never been analysed in a cell-free environment, i.e., without the influence of other cellular components. During these analyses, we noticed a particular dependence of TARA76 on the electrolyte used. More specifically, to deepen the proton pump functionality, we conducted experiments under two main conditions. The two chambers of the device were filled with buffered solutions either containing or lacking the Na^+^ ion, while the pH was varied or a different electrolyte was added. We would like to emphasize that all these experiments, carried out with various solutions, were performed in the absence of any protonophore or ionophore in the membrane (e.g., monensin or valinomycin), unlike what has been previously reported for this type of measurement.^44,48^ It was indeed shown that, apart from the transient current peak, a sustained stationary current is typically associated with the simultaneous presence of a proton pump and a passive transport channel. In our case, TARA76 within the membrane is the only bio-component expected to generate and sustain the photocurrents.

In the case of a Na^+^-containing buffer solution (10 mM NaCl and 20mM Na_2_HPO_4_/NaH_2_PO_4_), a clear stationary photocurrent was observed to increase when moving towards acidic conditions, particularly at pH 5 (Fig. 3a). This is consistent with enhanced proton pumping activity facilitated by the higher availability of free H^+^ ions. Across all tested pH levels, a transient peak was consistently observed upon light activation, suggesting an initial fast conformational switch or stronger proton pump activity.^43,44,48^ The amplitude of the stationary current exhibited also a further dependence on the identity of the surrounding cation (added to the buffer solution at pH 6, Fig. 3b). The presence of K^+^ and Ca^2+^ (Fig. 3b) produced a weaker response, 0.06 pA and 0.02 pA, respectively, compared to Na^+^ (Fig. 3b), which generated a stronger response of 0.35 pA. The marked dependence of the photocurrent on the type of cation supports the hypothesis that TARA76 could exhibit passive transport selectivity for Na^+^ ions,^55^ or that Na^+^ plays a structural role in stabilising the protein to preserve its functional activity. The photocurrent was then investigated using Tris/HCl, a different buffer without sodium. Despite a gradual increase in the stationary current when reducing the pH (Fig. 3c, also see Supplementary Fig. S6), once again an even larger current is achieved when the Tris-HCl buffer was used in combination with the presence of the Na^+^ ion (NaCl at 10 mM, Fig. 3c). We confirm in this way that the observed current is predominantly proton-based,^44,48^ yet supporting a fundamental role for Na^+^. On the other side, pushing even further the acidic conditions, leads to a reversal of the trend and a decrease of the stationary current. This is shown in the transients in Fig. 3d (also see Supplementary Fig. S7), by using an unbuffered solution (30 mM NaCl) without (magenta line) or with (black line) the simultaneous addition of 10 mM HCl (approximately equivalent to pH 3). In the latter case, the photocurrent decreases on a faster time scale. This is compatible with the strongly acidic environment leading to stable protonation of the residues involved in H^+^ transport, such as Asp78. The stable protonation is a mechanism known for driving proteorhodopsins into an inactive form,^26,43^ where the proton-pumping activity decreases rapidly or disappears completely.^43,56^

**Figure 3.**
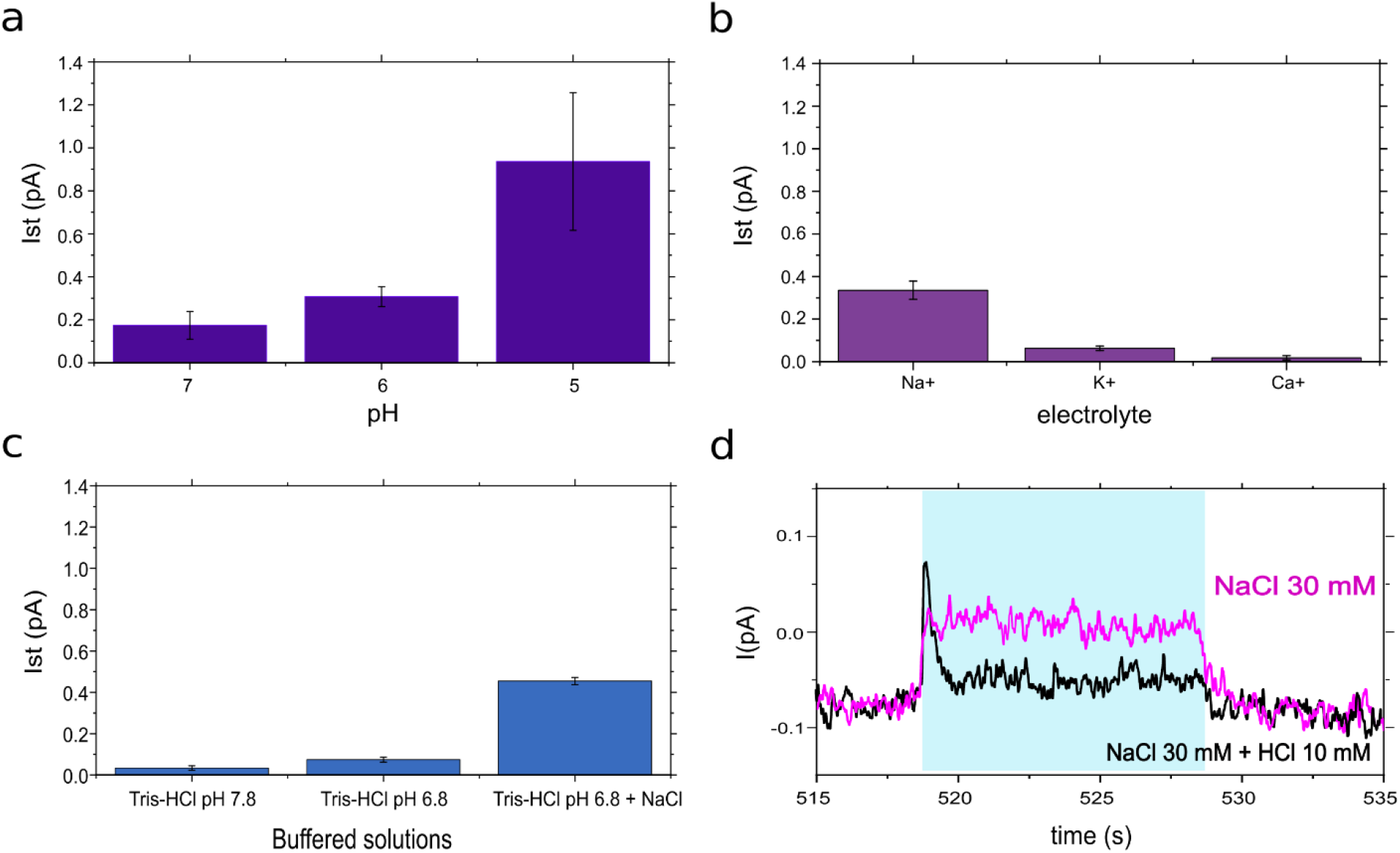
Dependence on pH and electrolye. (**a**) Histogram of light-induced stationary current extracted from the irradiation of TARA76-NDs incorporated into the lipid membrane at pH 7, 6, 5 in a buffered solution of 10 mM NaCl and 20 mM Na_2_HPO_4_/NaH_2_PO_4_. (**b**) Histogram obtained using three different buffered solutions at pH 6: (i) 10 mM NaCl + 20 mM Na_2_HPO_4_/NaH_2_PO_4_, (ii) 10 mM KCl + 20 mM Na_2_HPO_4_/NaH_2_PO_4_, (iii) 10 mM CaCl2 + 20 mM Na_2_HPO_4_/NaH_2_PO_4_. (**c**) Histogram corresponding to the following solutions: 20 mM TrisHCl pH7.6, 20 mM Tris/HCl pH 6.8, 20 mM Tris/HCl pH 6.8 + 10 mM NaCl. (**d**) Current tracings after adding TARA76-NDs in the upper chamber of the microfluidic device filled with the unbuffered solution of 30 mM NaCl (magenta curve, 10 s light exposure) and after the addition of 1 mM HCl (black curve, 10 s light exposure). The values of the stationary current in the histograms of panels a, b and c are obtained by averaging the current traces over 1000 ms interval of the stationary current; the error bars are estimated across repeated light exposure steps.

Taken together, these characterisations indicate that TARA76 primarily functions as a light-driven proton pump; however, its activity is dependent on the presence of Na^+^, which appears to play a key role in stabilising the protein in its functional conformation. Rather than acting solely as an external modulator, sodium significantly enhances the photocurrent amplitude by promoting or maintaining the correct structural folding of the protein, as supported by the TARA76 structural model simulations. In the absence of Na^+^, TARA76 show a less productive conformational state, resulting in reduced proton transport efficiency. This strong structural dependence on sodium raises the question of whether Na^+^ may also interact more directly with the transport mechanism, potentially through selective permeability. Indeed, several proteorhodopsins and channelrhodopsins have been shown to support dual Na^+^ /H^+^ fluxes, often with distinct efficiencies or kinetic profiles for each ion.^44,46,47,57,58^ Even though our data primarily support a structural and stabilising role for Na^+^, the possibility of a coupled Na^+^ /H^+^ transport mechanism remains an intriguing hypothesis that will require direct ion transport assays, site-directed mutagenesis, and high-resolution structural studies to be substantiated. The possibility that this cation helps protein stabilisation is corroborated in the following discussion in relation to modelling data.

### TARA76 structural model and the role of sodium

To provide a framework for the analysis of structure/function relationships of TARA76, a structural model has been built using the state-of-the-art ab initio protein structure prediction software AlphaFold 3.0 ^59^ and compared to the three-dimensional structure of the prototypical ion pump bacteriorhodopsin from *Halobacterium salinarum* (PDB code 1AP9 ).^60^ As can be seen from Fig. 4a, the TARA76 structural model is perfectly superimposable to the three-dimensional structure of BR. Further, all the amino acid residues involved in the proton-pumping mechanism in BR ^60^ have a counterpart in TARA76 (Fig. 4b), reinforcing the hypothesis that TARA76 is a bona fide light-driven proton pump.

**Figure 4.**
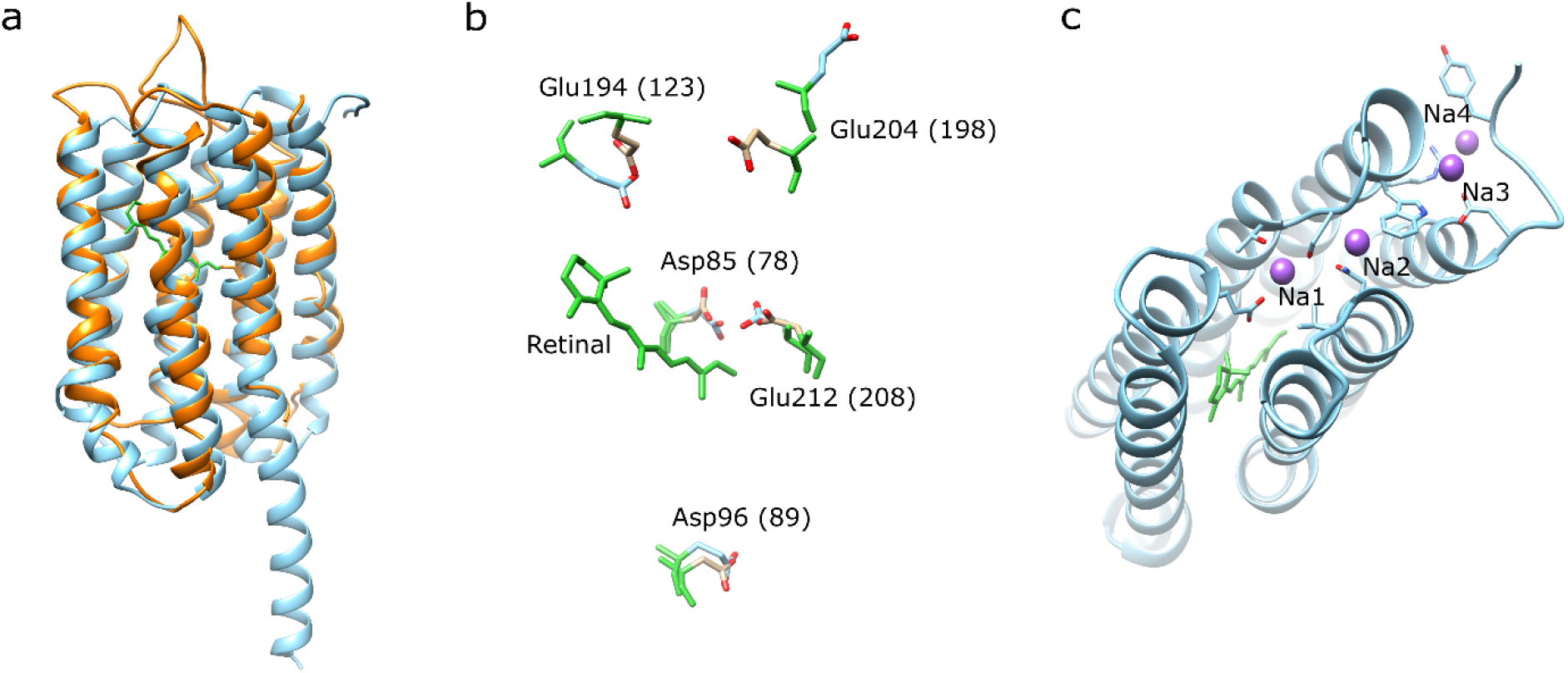
Details of the structural model of TARA76. (**a**) Structural superimposition between the crystal structure of *Halobacterium salinarum* bacteriorhodopsin (orange; PDB code 1AP9, Pebay-Peyroula et al., 1997) and TARA76 structural model (light blue). The retinal cofactor is shown in stick representation and coloured green. (**b**) Residues involved in proton conduction in bacteriorhodopsin (light brown) and orthologous residues in TARA76 (numbering shown in parentheses). For clarity, the residues’ backbone is coloured green. (**c**) Putative sodium binding sites of TARA76 predicted by AlphaFold 3.0.

The same modelling approach has also been used to predict putative Na^+^ binding sites in TARA76 to provide a structural basis for the effects of this cation observed in light-induced stationary current measurements. Indeed, AlphaFold 3.0 predicts the presence of several potential cation binding sites on TARA76 (Fig. 4c). Particularly interesting is the Na^+^ binding site indicated by the label Na1 in panel c. In this site, the cation is surrounded by several electronegative oxygen ligands, among which are those belonging to the carboxyl group of Glu123, which is predicted to be the orthologous residue of Glu194, a residue involved in the proton-pumping mechanism of BR.^60^ This may suggest the involvement of sodium in protein coordination, promoting the alignment of membrane proteins towards the same conformation. Sodium thus appears to selectively stabilise a specific conformation, leading to an increase in the current signal, as observed in Fig. 3b.

### AC analysis of TARA76 photoresponse

In previous works, the photoactivity of similar proteins in an artificial membrane has been studied by exclusively looking at the light-induced transient and stationary current in continuous-voltage, i.e., DC conditions. This has typically been associated with the conditions of ionic imbalance induced by the protonophore protein.^44,48^ To further investigate the vectorial nature of TARA76 photocurrent, we used an AC scheme. The AC is important because it can simultaneously look at active and passive components of the photoactivity; moreover, it helps prevent ion accumulation and excessive drift of electrical quantities. By analysing the dynamical response, it is also possible to study the capacitance of the membrane, potentially implicated in photo-induced conformational switching. We employed the same device and experimental setup, but we applied across the electrodes a triangular voltage waveform and registered the current response (Fig. 5a, dashed black trace and solid blue trace, respectively).

**Figure 5.**
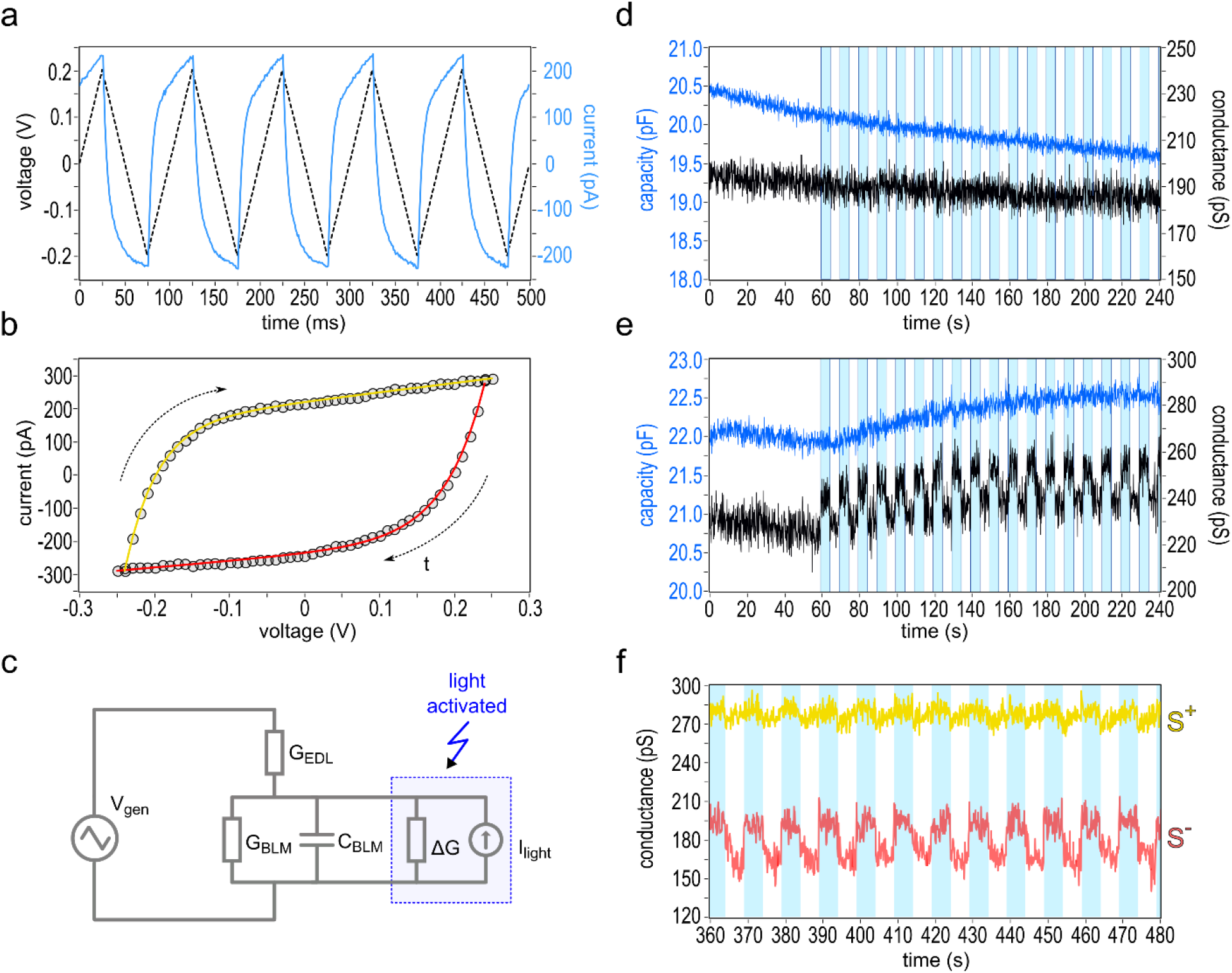
AC experiments. (**a**) Time charts of the AC triangular voltage (dashed red line) and the modulated current output (solid blue curve) along five periods (modulating amplitude V_m_ = 100 mV, frequency 10 Hz, sampling time 1 ms). (**b**) The I(V) cycle over a single 100 ms period at a modulating amplitude V_m_ = 250 mV (grey circles). The fit of the two half-wave branches is superimposed (solid yellow and red lines). (**c**) Reduced equivalent circuit for a planar BLM bilayer plus a series EDL. (**d**) Conductance (black line) and capacitance (blue line) time traces extracted from the fit of the current waveform in the case of empty NDs added to the membrane. The laser exposure windows are indicated by shaded blue area (5 s light-on each followed by 5 s light-off intervals). (**e**) Conductance (black line) and capacitance (blue line) with TARA76-loaded NDs in the membrane. (**f**) Positive (yellow line, S^+^) and negative (red line, S^-^) conductance from the fit of the two half-cycles, in the same case of (e). The time step in panels (a,b) is 1 ms, in panels (d,e) is 100 ms.

In Fig. 5b, we show a typical current-voltage or I(V) cycle at a modulating amplitude V_m_ = 250 mV. In the absence of the membrane, the I(V) curves are practically linear, according to the highly conductive solution. Instead, following the membrane formation, the I(V) curves assume a hysteresis-loop shape, indicating a predominant capacitive behaviour. To extract the capacitive and resistive components, we used a fitting model able to reproduce the experimental results. This is associated with the reduced equivalent circuit reported in Fig. 5c (see Supplementary Fig. S8a for the complete circuit ).^56,61–63^ The resulting current under a triangular voltage waveform can be written as I(t) = G_tot_*V(t) + C_BLM_*(dV/dt) + T*exp(-t/τ), where I is the measured current and dV/dt is the time derivative of the triangular voltage applied by the generator. This equation provides a linear term for each half-wave branch, with a slope equivalent to the series conductance G_tot_ = 1/[(1/G_BLM_) + (1/G_EDL_)] ≈ G_BLM_ (where the BLM term is dominant), and a vertical shift proportional to the capacitance C_BLM_, plus an exponential T term with time constant τ. The first two terms are sufficient to reproduce the merely parallel RC element of a leaky BLM membrane. However, in this case, the hysteresis loop would be a parallelogram (see Supplementary Fig. S8b, c). Instead, to reproduce the round edges of the cycle in the second and fourth quadrants of the IV plane, an appropriate resistive element is needed in series, here G_EDL_. This represents the two electric double layers (EDL)^56,63^ at the membrane-solution interfaces. Its exponential response is defined by a time constant τ ∼ 5 ms. Finally, the ΔG and I_light_ elements represent a non-ideal current generator associated with TARA76 photo-stimulation, consisting of a parallel conductance variation and a current source.

The capacity and conductance are obtained as averages from the separated fits of the two half-wave branches. Their behaviour over time is shown for the artificial membrane incorporating either empty nanodiscs or TARA76-loaded nanodiscs in Fig. 5d and 5e, respectively. In accordance with the DC experiments, no light-dependent variation of conductance is observed in the absence of TARA76 (Fig. 5d, black line). Oppositely, light-induced conductance steps are neatly visible in the presence of TARA76 (Fig. 5e, black line). These steps are in the order of ΔG = 20 pS and almost instantaneous on the time scale of the AC experiments (where each fit is performed every 100 ms). Moreover, the conductance steps are positive, which aligns with the positive photoconductance seen in the DC transients (where ΔG = 1 pS was associated with the I_st_(V_DC_) slope). The AC photoconductance exceeds the DC by an order of magnitude, due to the larger bandwidth associated with the smaller amplification factor required in the AC case. Differently, the extracted capacitance shows no appreciable modulation during the light exposures, both in the case of empty NDs (Fig. 5d, blue line) and in the presence of TARA76-NDs (Fig. 5e, blue line). A slow and slight (1%) increase of the capacitance was observed under cyclic irradiation in the presence of TARA76, whose plateau suggests a possible light-induced conformational stabilization. The same but larger (10%) light-induced slow drift of G may indicate instead that the continuous proteins’ activity gradually alters the EDL at the two membrane-solution interfaces. Similar behaviour of both C and G has been observed in repeated experiments at different V_m_ (Supplementary Figs. S9-S10 and their discussion).

### Vectorial nature of TARA76 photoactivity

We note that the triangular AC allows to finely investigate for a potential asymmetry in the electrical and photoresponse. From the separate fit of the two half-wave branches of the loop (yellow and red thick solid lines in Fig. 5b), we retrieve the conductance along both the positive and negative directions of the voltage scan. These are labelled as S^+^ (yellow line) and S^-^ (red line) in Fig. 5f, respectively, where are shown during several time intervals of light-on/-off. Interestingly, a much higher light-induced modulation of S^-^ is observed than S^+^. In general, this asymmetric behaviour suggests a preferential orientation of the protein within the membrane. It is important to note that the complete AC analysis allows in principle to assess the ongoing TARA76 photoactivity and its vectorial nature, also when there is no visible modulation of the average current. The observed behaviour is compatible with a light-activated electrogenic process with both conductive and active transport components (for more details see Supplementary Fig. S11). In our case, we think that the microscopic directionality of the protein may be extended to the whole membrane by a natural mechanism. When injecting the NDs only from one side, a different affinity of the membrane for one side of the proteins can increase the preferential orientation of TARA76-NDs incorporation. However, this natural affinity may be weak in general and, since TARA76 is reconstituted in nanodiscs and not injected by means of functionalised liposome, controlling its orientation may be challenging. Strikingly, we have also found an orientational forcing mechanism, upon applying a DC voltage during the membrane formation itself, in the presence of TARA76. Similarly to electric poling treatment for organic molecules, we can induce a preferential orientation, therefore controlling the resultant photocurrent direction (Supplementary Figs. S12-13). This is a very important asset in the engineering of photo-responsive electrical features of either protein pumps or channels embedded in a bio-mimetic membrane.

## Conclusions

In this work, we studied the photoelectrical behaviour of the recently discovered microbial rhodopsin TARA76 in a fully artificial, acellular membrane system, revealing its potential as an active component for bio-photonic devices. By reconstituting TARA76 into nanodiscs and integrating them within a black lipid membrane, we succeeded in recording reproducible photocurrents under controlled optical and electrical conditions, isolating the intrinsic properties of the protein from cellular interference.

Importantly, we demonstrated that the protein’s response can be modulated by both the ionic environment and the applied voltage, unveiling a marked dependence on Na^+^ ions not previously reported for this class of proteorhodopsins. This observation broadens TARA76’s functional scope beyond that of a simple proton pump. We further showed that the orientation of the protein within the artificial membrane can be externally guided by an applied field during bilayer formation, offering a practical route to control the directionality of the photocurrent, a crucial step toward engineering bio-electronic interfaces with defined polarity. Moreover, the light-induced modulation of conductance observed under AC excitation provides additional evidence of asymmetric conduction confirming the vectorial nature of its light-driven current-generator activity and its extension to the whole membrane. Our results provide the first detailed analysis of TARA76’s photocurrent shape, kinetics, and amplitude, identifying its activation and relaxation dynamics.

Overall, these results establish TARA76 as a robust and stable photoactive protein capable of operating at room temperature. The acellular micro-device we developed constitutes a versatile element for next-generation bio-photon detectors, neuromorphic components, and bio-hybrid optoelectronic systems. TARA76 is therefore a perfect candidate for future studies of bio-compatible platforms to be embedded into compact, low-noise, and wavelength-selective detectors as well as biomedical applications such as artificial retinal prostheses.

## Material and methods

### Microfluidic device features and artificial membrane formation

The disposable microfluidic device consisted of two compartments of approximately 150 μl, separated by a PTFE septum containing a single microscale aperture that connects the two chambers. Each chamber included one compartment for immersed Ag/AgCl electrodes. They were plugged into the head stage’s BNC port and the ground port at the side of the head stage. The artificial membrane was produced using the air bubble technique. A drop of 10mg/ml of DPhPC (1-palmitoyl-2-[3-(4-((1E,3E,5E)-6-phenylhexa-1,3,5-trien-1-yl)phenyl)propanoyl]-sn-glycero-3-phosphocholine, Avanti® Polar Lipids) solution in n-octane (Sigma Aldrich at 99% purity used as received) was pipetted in proximity to the microaperture in a Teflon septum. Two device types were used: Ionovation (Ionovation GmbH, Germany) and Elements (Elements srl, Italy), with nominal aperture diameters of ∼120 µm and ∼80 µm, respectively. After formation, the BLM set horizontally between two microchambers filled with the electrolyte solution (0.5 M KCl, 20 phosphate buffer pH 7.6 provided by ChemCruz and solubilised in Milli-Q® ultrapure H_2_O), each independently accessible and provided by electrodes. The lipid phase between the two water compartments underwent a spontaneous, energy-minimizing thinning process that ultimately yielded a single-bilayer membrane. This formation was monitored optically and by capacitance measurements. These lasts were performed by measuring the electric current between the chambers upon applying a triangle wave voltage (amplitude 100 mV, frequency 10 Hz). The Ag electrodes were chlorinated, immersing them in a bleach solution. Afterwards, the Ag/AgCl electrodes were directly immersed in the electrolyte solution through two holes connected to each chamber and simultaneously to the current amplifier’s head stage (Low Noise Current Preamplifier, Standford Research System). The amplified currents were digitised at a sampling interval of 1 ms and fed into a converter (National Instrument, eight inputs, 16-bit, 2.0 MS/s X Series Multifunction DAQ) for storage on a personal computer. A custom software in the language NI-LabView was used to interface instruments and monitor the membrane formation and its capacity in real-time during experiments. Same environment was also used together with OriginPro 2018 (Microcal Software Inc.) to analyse data.

### DNA cloning

Dr. Francesco Calabi performed the cloning experiments and produced every expression vector described below at CNR-Nanotec. The coding sequences for the *Escherichia coli* expression of TARA76 were selected from Shim’s works.^28,44^. Eurofins Scientific synthesised the DNA. The genes were optimised and introduced in a pET(28a) (provided by Addgene) expression vector using two restriction sites. An additional His-tag sequence was added to the C-Terminus of the protein-coding sequence.

### Nanodiscs and liposome formation

For Nanodiscs formation, 1,2-dimyristoyl-sn-glycero-3-phosphocholine (DMPC – Avanti Polar Lipid) stock was prepared by suspending lipids in Milli-Q® H_2_O to a final concentration of 100 mM. For each lipid, a particular final concentration of sodium cholate (ChemCruz) may be required to obtain an apparent suspension, in this case, 200 mM of Sodium cholate. The MSPE3D1 scaffold proteins were expressed in *E. coli* BL21 (DE3) transformed with Addgene plasmid 20066 as previously described ^33^. This protein had N-terminal poly-histidine tags used for its purification using affinity chromatography with HisPur™ Ni-NTA resin column (EMD Millipore corp, Merk). All the collected fractions were concentrated by an Amicon Ultra-4 3K centrifugal filter tube (Millipore Corp, Merk) and kept at 4 °C until use. Concentrations were determined from the absorbance at 280 nm using calculated extinction coefficients. ^64^ All buffers consisted of 10 mM Tris–HCl, pH 7.4, 0.1 M NaCl. For ND assembly ^31,33,52,65^ The MSP1E3D1 was mixed with DMPC and cholate at a molar ratio of 1:80 and incubated for > 2 h at 24 °C, followed by detergent removal with an XAD-2 resin. Amberlite XAD-2 was obtained from Sigma Aldrich. The Amberlite beads were washed three times with methanol (Sigma Aldrich) and then equilibrated and stored in distilled water. The amount of Xad for the detergent removal is calculated based on the detergent micromoles and was calculated experimentally as described previously.^66,67^. For cholate removal, the ratio of mg XAD/mmol cholate is 1:8 approximatively.

### Expression and purification of TARA76

The protein was expressed in *E. coli* BL21 (DE3). After transformation, cells were grown in Luria–Bertani (LB) medium supplemented with kanamycin (50 mg/mL; Sigma-Aldrich) at 37 °C. An overnight culture was used to inoculate 500 mL of LB medium containing kanamycin (50 mg/mL), and the culture was incubated at 37 °C with shaking at 300 rpm until it reached an optical density at 600 nm (OD_600_) of 0.4–0.6, measured using a Cary50 spectrophotometer (Cary WinUV Pharma). Protein expression was induced by adding 1 mM IPTG and 7 μM all-trans-retinal (Sigma-Aldrich), followed by incubation for 6 h at 37 °C. Cells were harvested by centrifugation at 5,000 × g for 15 min (Eppendorf centrifuge 5810R), washed with buffer (150 mM NaCl, 50 mM Tris-HCl, pH 7.0), and centrifuged again at 4,000 × g for 20 min. The resulting pellet was resuspended in lysis buffer and homogenized three times for 10 min each. The samples were then ultracentrifuged at 20,000 × g for 15 min. The pellet was resuspended in lysis buffer supplemented with 2% LDAO (N,N-dimethyl dodecylamine N-oxide; Sigma-Aldrich) and 10 μM all-trans-retinal, and solubilized overnight at 4 °C. The suspension was subsequently ultracentrifuged at 20,000 × g for 30 min. The supernatant was diluted to a final concentration of 0.2% LDAO and supplemented with 20 mM imidazole. It was then incubated with Ni^2+^-NTA agarose resin (Qiagen) under gentle shaking for 2 h at 4 °C. Following this step, purification was carried out as previously described. Finally, the protein was concentrated using an Amicon Ultra-4 10 kDa centrifugal filter unit and stored at 4 °C until use.^28,44^

### Reconstitution of the protein into nanodiscs

For ND –IMP reconstitution reactions, we followed the previous protocols ^31,53,54^ with some modification given by the size and the tertiary structure of the proteins that must be integrated. The MSP1E3D1 was mixed with DMPC/cholate and the rhodopsins at a molar ratio that was respectively 1:126.5:0.5 (reaction A) and 145:1:0.1 (reaction B) for TARA76 and 1:100:0.25 ^54^ For ChRmine. They were incubated for 30 minutes at 24 °C without the MSP and 12 h with the MSP. This incubation was followed by detergent removal with XAD-2 resin (Amberlite, Sigma Aldrich). For detergent removal, the presence of LDAO was also considered. The resulting nanodiscs were analysed by absorbance spectroscopy to evaluate TARA76 and ChRmine functional characteristics. The empty nanodiscs used as a control were produced by adapting the preview protocols.^31,32,45,53^ The illustration of NDs reaction in Figure 1 was performed using © 2026 BioRender (www.biorender.com, Scientific Image and Illustration Software).

### Biochemical proteins characterization

Protein samples were analyzed by denaturing sodium dodecyl sulfate–polyacrylamide gel electrophoresis (SDS-PAGE). Samples were mixed with Laemmli sample buffer containing SDS (Sigma Aldrich) and a reducing agent (either β-mercaptoethanol, Sigma Aldrich) and heated at 95 °C for 5 minutes to ensure complete denaturation. Proteins were separated on polyacrylamide gels (typically 12% resolving gel) prepared according to standard protocols. A stacking gel (4% acrylamide) was used to improve band resolution. Electrophoresis was carried out in running buffer (25 mM Tris, 192 mM glycine, 0.1% SDS) at 70 V for the first 30 min and 120 V until the end. A prestained SDS-PAGE protein marker broad range (Bio-Rad) was loaded alongside samples to estimate protein sizes. Following electrophoresis, gels were stained with Coomassie Brilliant Blue R-250 using an acid-based protocol. Briefly, gels were incubated in staining solution containing 0.1% Coomassie Brilliant Blue R-250 dissolved in aqueous hydrochloric acid (0.1 M HCl) at elevated temperature (typically 70 °C) for 30 min with gentle agitation. After staining, excess dye was removed by 3 washes of 5 min each in distilled water at the same temperature until a clear background and well-defined protein bands were obtained. Gels were subsequently imaged using a ChemiDoc imaging system (Bio-Rad), and images were processed using the Bio-rad software.

### Spectroscopic characterisation of Chronos, ChRmine, TARA76 and OLPVRII

We recorded the absorption spectra using the Nanodrop one (Thermofisher) spectrophotometer. The spectrum for each protein was determined in a range between 200 nm to 900 nm and was taken in the dark at room temperature. The measurements were then analysed with OriginPro 2018.

### Dynamic light scattering

The measurements were performed on a Zetasizer nano series (Malvern Instruments) at 25 °C in disposable 50 μl micro cuvettes. Data were collected for 20 s; an average of 15 scans were performed for liposomes, and 25 scans were performed for nanodiscs. The size distribution is shown on a log scale using the OriginPro2018 software.

### Transmission Electron Microscopy (TEM)

An empty-ND sample solution, diluted to 10 ng/ml, was drop casted onto a standard carbon coated Cu TEM grid, allowing solvent to evaporate. The as dried sample grid was imaged by using a JEM 1400Plus (JEOL Ltd. – Japan) microscope, operating at 120 kV with a LaB_6_ source.

### Photoelectric measurements and optoelectronic setup

The optical part consists of an optical microscope (Olympus 1x2-UCB) through which it can visualise and excite the membrane. Using an optical path on an optical table, the microscope image is projected onto the entrance slits of an imaging spectrometer (Horiba, iHR320) equipped with a 300 lines/mm grating and coupled to a 2D enhanced charge-coupled device (Hamamatsu, EMCCD C9100). The membrane is digitalised on a computer and visualised using Windows-supported software (Hamamatsu, HPD-TA High-performance Digital Temporal Analyzer – 64bit. At the same time, it is possible to get the imaging of the membrane (by using a halogen broadband white light source in a transmission configuration) and carefully align the laser (reflection configuration) to direct the beam in the membrane centre. A lateral aperture in the microscope allows the laser to be coupled and aligned thanks to a series of mirrors and lenses (ThorLabs Inc.). The laser beam is then focused onto the membrane using a 20X objective (Olympus) in a spot size with a Full Width at Half Maximum (FWHM) comparable to the hole in which the membrane is formed. We used different types of lasers: Continuous Wave (CW) Diode Lasers at 488 nm and 532 nm (MKS-SpectraPhysics) and a tunable pulsed laser (Computer-Controlled Femtosecond Optical Parametric Amplifier, TOPAS-prime, Coherent) set at different wavelengths (480 nm, 490 nm, 500 nm, 510 nm, 520 nm, 530 nm, 540 nm, 550 nm, see Supplementary Fig. S4 for more details). The opening and closing time of the laser was controlled by using an optical shutter (Thorlabs, SH05R) interfaced with a computer and controlled by a custom NI-LabView program, as described in the paragraph above. This made possible to control and synchronize the applied voltage waveforms and laser exposure, according to different protocols, such as changing either a V_DC_ or V_m_ in a sequence, for example going from -125 mV to 125 mV in steps of 25 mV and applying cyclic irradiation time-windows for each point.

## Acknowledgements

The authors gratefully thank Franco Calabi for helpful discussions and mentorship, for helping in the idea conception and work planning; Ilaria Palamà, Gabriele Maiorano, and Gabriella Leccese for their scientific support; and Paolo Cazzato, Sonia Carallo, Laura Polimeno and Vincenzo Mastrolia for their valuable technical assistance and scientific support. This work was supported by the project “Tecnopolo per la medicina di precisione” (TecnoMed Puglia)—Regione Puglia: DGR n.2117 del 21/11/2018, CUP: B84I18000540002, by the European Union - NextGeneration EU, PNRR MUR “Integrated infrastructure initiative in Photonic and Quantum Sciences” - I-PHOQS [IR0000016, ID D2B8D520, CUP B53C22001750006], by the PNRR MUR National and Quantum Science Technology (NQSTI) – Spoke 2 (CUP B53C22004180005, PE0000023), by the project ‘Quantum Optical Networks based on Exciton-polaritons (Q-ONE)’ funded by the HORIZON-EIC-2022-PATHFINDER CHALLENGES EU programme under grant agreement No. 101115575 and by the project ‘Neuromorphic Polariton Accelerator (PolArt)’ funded by the Horizon-EIC-2023-Pathfinder Open EU programme under grant agreement No. 101130304. L.D.M. acknowledges the European Union - NextGeneration EU project “Network 4 Energy Sustainable Transition – NEST” (Project code PE0000021, CUP B53C22004060006, Concession Decree No. 1561 of 11.10.2022 adopted by Ministero dell’Università e della Ricerca) and the ERC Consolidator project no. 101045746 — HYNANOSTORE. C.N. acknowledges the project “Fit for Medical Robotics”-Fit4MedRob Grant (Code: PNC0000007, CUP: B53C22006960001). Views and opinions expressed are however those of the author(s) only and do not necessarily reflect those of the European Union. Neither the European Union nor the granting authority can be held responsible for them.

## Author contributions

D.S. conceived the idea, together with Franco Calabi to whom D.S., I.C., L.D. and L.D.M. are grateful for his fundamental help and support. D.S. and L.D.M. coordinated the experimental work. I.C. performed the protein expression, extraction, and purification; the biochemical characterizations, the spectroscopic analysis in solution; the optimization of artificial membrane formation; and the photoelectrical measurements on the microfluidic devices. C.N. performed TEM measurements and analysis. A.C. built the electrical setup and L.D. developed the software program for interfacing and analysing the photoelectrical experiments. L.D., A.C., V.A., D.S. and L.D.M supervised the photoelectrical measurements on the microfluidic device. V.A., A.F., L.D., D.B. and M.D.G implemented the optical setup. V.A., A.F. and D.S. supervised the optical measurements. I.C. and L.D. analysed the photoelectrical data. F.P. performed the structural modelling and with F.B. analysed the conformational behaviour of TARA76. I.C. wrote the first draft of the manuscript. L.D., L.D.M. and D.S. carefully reviewed the original draft. All authors contributed to the discussion and interpretation of the data and revised the manuscript.

