## Supplemental Information for "Electrically programmable picoscale phototransduction of a newly discovered microbial rhodopsin"

---

*[1]CNR-NANOTEC, Institute of Nanotechnology, via Monteroni, 73100 Lecce, Italy; [2]CNR-IMM, Institute for Microelectronics and Microsystems, via Monteroni, 73100 Lecce, Italy; [3]Department of Science, University of Roma Tre, 00146 Roma, Italy; [4]Department of Experimental Medicine, University of Salento, c/o Campus Ecotekne, via Monteroni, Lecce, Italy.*

***Sequence alignment.*** TARA76 is a protein that was isolated from biological samples collected by the *Tara* Oceans expedition. Its entire sequence was recently obtained, and the protein was expressed in bacteria, which were characterised in different extreme conditions.[1] Sequence alignment of TARA76 rhodopsin with different microbial rhodopsins confirmed a remarkable similarity with the proteorhodopsin group.[1] TARA76 absorbs light in the 480-530 nm range, depending on the pH. To understand the structure and the possible characteristics of TARA76, the sequence alignment was here performed with a BPR (blue-light-absorbing proteorhodopsin), a GPR (green-light-absorbing proteorhodopsin) and *Med12*BPR (a BPR isolated from the Mediterranean Sea at a depth of 12 m), proteins that perform ion-pumping functionality like the archaeal homolog bacteriorhodopsin. *Med12*BPR also has the defined crystallographic structure and significant homology with TARA76 and the other two proteins. This alignment shows that the DTE motif associated to the proton pump is present in all four PRs, as well as the L/Q105 which is known as the colour-tuning residue. According to Ran and coworkers[2], as mentioned before, the light-driven ion-translocation process is characterised by two crucial residues, Asp97 and Glu108 (green square, Supplementary Fig. S1a), that have been demonstrated to act as the primary Schiff-base proton acceptor and donor, respectively, in *Med12*BPR. Both are present in the TARA76 sequence. Also, the His76 (purple square, Supplementary Fig. S1a) is a conserved amino acid in all the proteorhodopsin. The sequence shows a high similarity percentage between transmembrane sequences between TARA76 and *Med12*BPR (bright green square). Besides the demonstration that TARA76 belongs to the proteorhodopsin family, the most exciting result of these measurements is the determination of the possible quaternary structure of TARA76. So, we focused on the amino acids that may be involved in the protomers interaction. The oligomerisation states of the two variants of PR differ not only from each other (hexamer versus pentamer) but also from that of BR (trimer). Analysis comparing the three assemblies reveals that while there is some overlap between the interfaces, there are apparent differences between BR and the PRs and even between the three PR variants (Supplementary Fig. S1b). Anyway, the sequences reveal that all the amino acids involved in the protomer interaction in the *Med12* structure were aligned with the other two proteorhodopsins and TARA76 (underlined in green). The interface regions involve different helices. In all the variants of PR, helix A is heavily involved in the interface, along with helix C. *Med12* and TARA76 have many interface regions in common, even in the N-terminal region of the protein in which there is almost no similarity between the other two PRs that have pentameric oligomeric structure. The most critical residues are, as mentioned above, His75(57)-Trp34(16)-Asp97(79)-Val53, and as shown in the alignment, all of them are present in TARA76 except for the V53 that is substituted with Ile53. The essential salt bridge R51-D52’ would be, therefore, less favourable since the additional Ser changes the relative side chain orientations of Arg51(33) and Asp52(35) in red (Supplementary Fig. S1b), which, however, could be compensated by including a sixth protomer into the ring-shaped complex. This suggests that TARA76 could have a hexameric structure like *Med12*. The alignment with bacteriorhodopsin has demonstrated the presence of the amino acids involved in the proton transport, underlining that TARA76 is a light-driven proton pump.

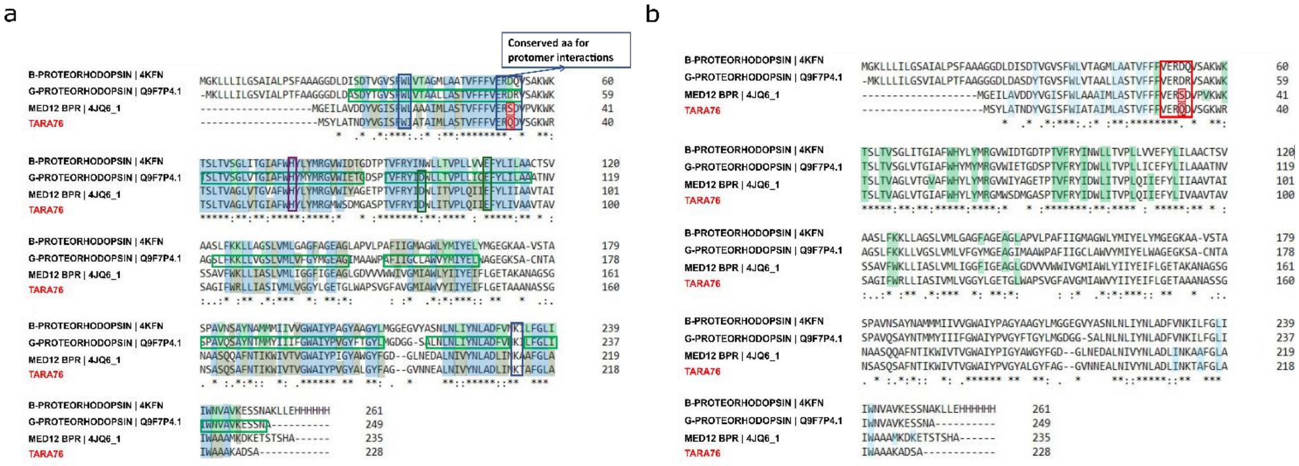


**Supplementary Figure S1. Sequence alignment analysis.** (**a**) Protein sequence alignment of different microbial rhodopsins: blue-proteorhodopsin 4KNF, green-proteorhodopsin Q9F7P4.1, *Med12*BPR and TARA76. The all-sequence similarity, which is 76%, is represented in blue, more precisely, 100% Similarity in the blue shaded area, 75% grey shaded area and 50% grey and green. Transmembrane regions are denoted by green squares above the alignment, which is 100% like *Med12*BPR and TARA76. Schiff base lysine (K) and all the conserved residues for the proton transport are annotated in the blue and green (amino acids D and E) empty squares above the alignment. His76 is a conserved amino acid in all the proteorhodopsins, and a purple empty square indicates it. (**b**) Alignment of the regions involved in oligomerisation. Residues denoted in blue are found on protomer A of each molecule and are involved in the interface with protomer B, represented in green. The residues in the hydrogen bonds or salt bridges are represented in red.

***TARA76 fingerprints and characterization.***

TARA76 was monitored with spectral absorbance, Dynamic Light Scattering (DLS), and SDS-PAGE (Sodium Dodecyl Sulphate - PolyAcrylamide Gel Electrophoresis) as discussed in the main text and in the caption to Supplementary Fig. S2.

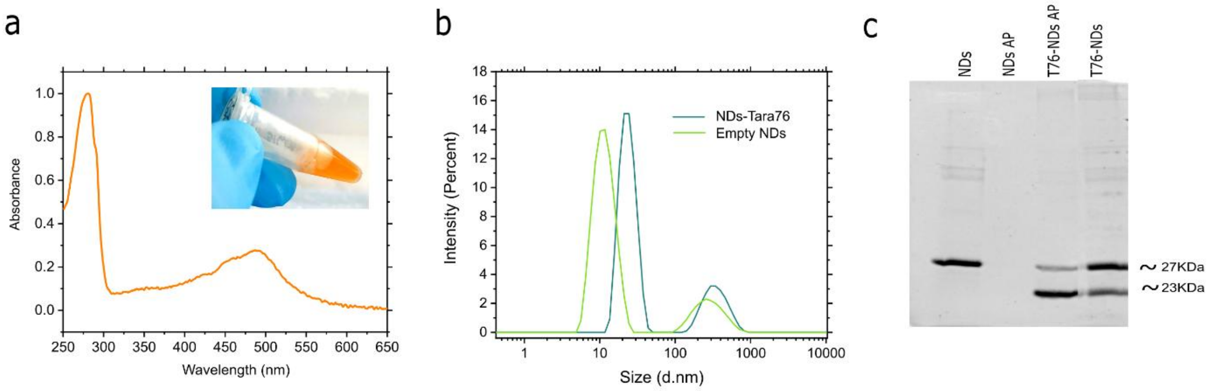


**Supplementary Figure S2** (**a**) Functional absorption spectra of TARA76 proterhodopsin (distinctive peak at 490 nm) reconstituted in nanodiscs with the reaction 1:126.5:0.5 ratio respectively for MSP: DMPC: TARA76 monomer. (**b**) Dynamic light scattering for TARA76-loaded NDs (dark green) and empty NDs (light green) diluted in a 10 mM NaCl phosphate buffer solution. The analysis was performed using a 50 µL disposable cuvette. (**c**) The nanodiscs assembly reaction was performed with TARA76-H6 and MSP1E3D1 without the histidine Tag(H6), and the reaction products were submitted to an IMAC purification process with Ni-NTA agarose resin. TARA76-loaded NDs and empty NDs were analysed with SDS-PAGE before and after the purification.

***Single channel measurements: Gramicidin.*** The starting measurements were performed with Gramicidin A to test the system sensitivity. These were conducted in DC mode, maintaining the constant voltage at 100 mV and measuring the current as an output signal. First, the BLM was tested for its stability without any addition. As shown in Supplementary Fig. S3a, the current is stable, and there is no change for at least 5 minutes. Following, Supplementary Fig. S3b shows the effect after the addition of Gramicidin A pore into the upper and lower chambers of the microfluidic device with a concentration of 50 ng/mL. Initially, the current recordings at a constant voltage of 100 mV show stepwise changes of ~ 5 pA consistent with the opening and closing of single Gramicidin A channels.

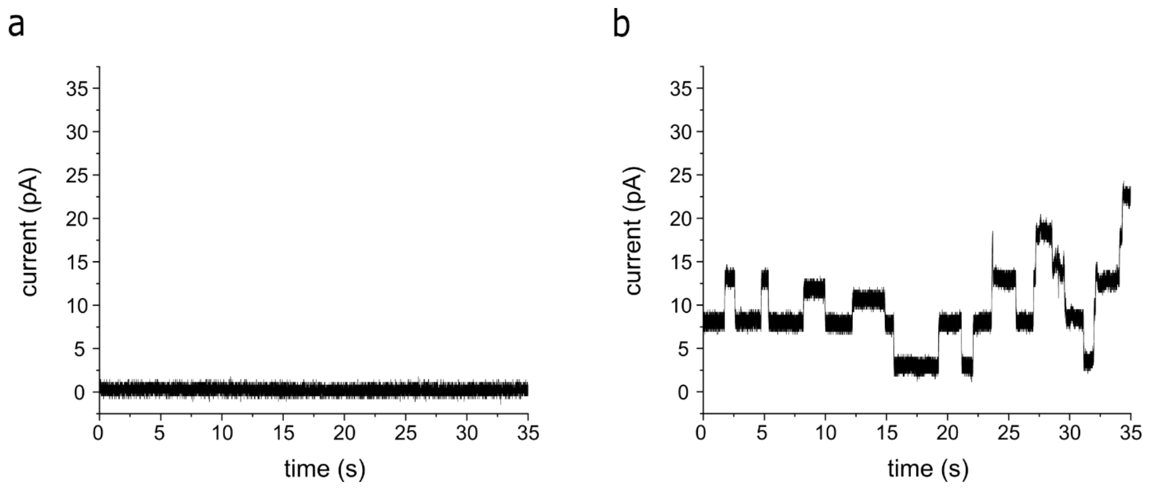


**Supplementary Figure S3** (**a**) Current tracing upon a 100 mV DC voltage of the BLM alone, (**b**) Single channel Gramicidin A (50 ng/μL) current measurement after its incorporation on the artificial membrane (BLM) formed in the Ionovation flow cell under 100 mV DC voltage.

***TARA76's current dependence on the spectroscopical characteristics of the protein.*** A spectral photocurrent measurement was performed irradiating the functionalised membrane with different wavelengths and measured the photocurrent per each. The protein was added to the microfluidic device at the maximum concentration, then the photocurrents were registered in succession and under the same conditions. The irradiation was performed with a pulsed laser (10 kHz of frequency) that was directed into the microscope. Different wavelengths were employed, starting at 480 nm and continuing until 550 nm, in steps of 10 nm. We can observe from the graph (Supplementary Fig. S4a) that there is a decrease in the photocurrent peak in dependence on the wavelength used. This is even more visible considering the stationary current (Supplementary Fig. S4b). It is observed that the trend is not linear, but the first 4 points of the graph have almost the same values, then at 530 nm, start to decrease until the achievement of 0 at 550 nm. Hence, the measured photocurrent is comparable to the functional absorption spectrum of TARA76, in which the absorption peak is at 490 nm at pH6. We could explore this part of the absorption spectrum and not shorter wavelengths, because the pulsed laser could damage the membrane and the device even at low intensity when used below of 490 nm.

For this reason, we compared the photocurrent given by the irradiation with two diode lasers at 405 nm and 480 nm, using a continuous wave irradiation instead of a pulsed one. The photocurrent measured irradiating with a 405 nm laser is lower than the ones at 488 nm (Supplementary Fig. S4 c,d) and 530 nm (pulsed laser, Supplementary Fig. S4e) indicating that it follows the absorption spectrum trend in the UV spectrum range that was impossible to observe with the pulsed laser. These spectroscopic results confirm that the photocurrent presented in the main text depends on the protein photo-absorption, and any other side effect can be excluded. [3,4]

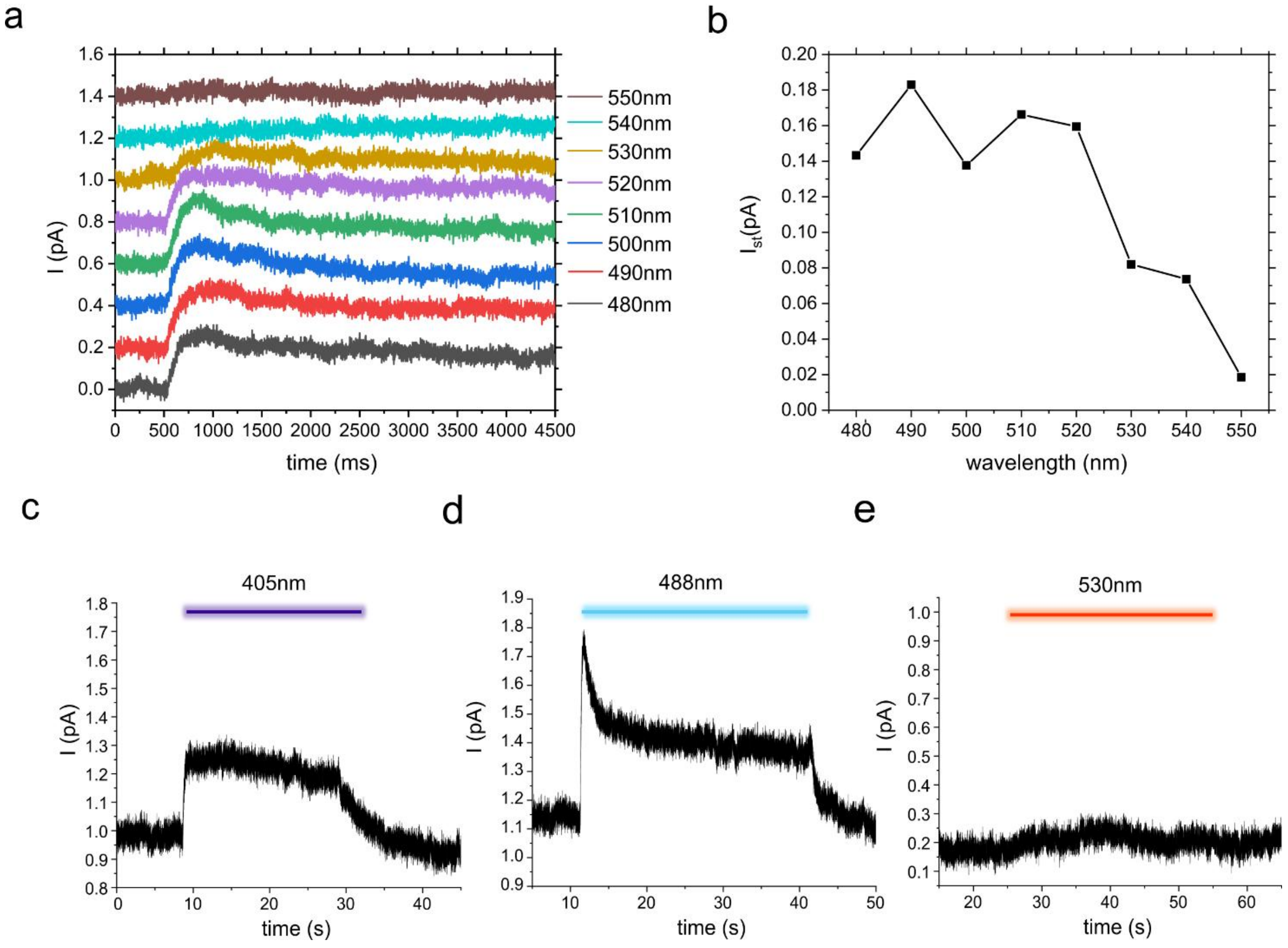


**Supplementary Figure S4** (**a**) Current trace recorded after incorporation of TARA76 expressed in *E. coli* (BL21) reconstituted in nanodiscs into a BLM. The membrane was irradiated using Topa's pulsed laser (10 KHz) at 40 μW of power, with wavelengths ranging from 480 nm to 550 nm in 10 nm increments. Measurements were conducted at pH6 in phosphate-sodium buffer under a constant voltage of 100 mV. (**b**) Wavelength dependence of the stationary photocurrent $I_{st}(\lambda)$.(**c-e**) Current transients recorded after incorporation of TARA76-NDs and under irradiation at specific wavelengths: 405 nm (c) and 488 nm (d) diode laser at 1 mW, and a 530 nm (e) Topa's pulsed laser (10 KHz repetition rate) at 40 μW.

***TARA76's current dependence on the concentration.*** A series of TARA76 dilutions was prepared, as detailed in the Experimental section. The resulting photocurrent responses at four concentrations, 0.53 mM, 1.075 mM, 2.15 mM, and 4.3 mM, are shown in Supplementary Fig. S5a. At each concentration, the primary photoresponse consisted of a rapid upward current increase under 1 mW laser exposure. Supplementary Fig. S5b illustrates the photocurrent as a function of TARA76-ND concentration. Notably, the response curve is nonlinear, indicating saturation of the signal at higher concentrations. Beyond a certain threshold, further protein addition compromises membrane stability, suggesting a maximum usable concentration before the integrity of the black lipid membrane (BLM) is disrupted. It should be noted that the acquisitions in Supplementary Fig. S5 are obtained using a lock-in amplifier. This system has a slow response in time since it selectively integrates over several time periods the signal oscillating at the reference frequency of the optical chopper (here 12 Hz). It does not capture rapid transient events, such as the initial burst in photocurrent upon light exposure, and its response is not rescaled in intensity.

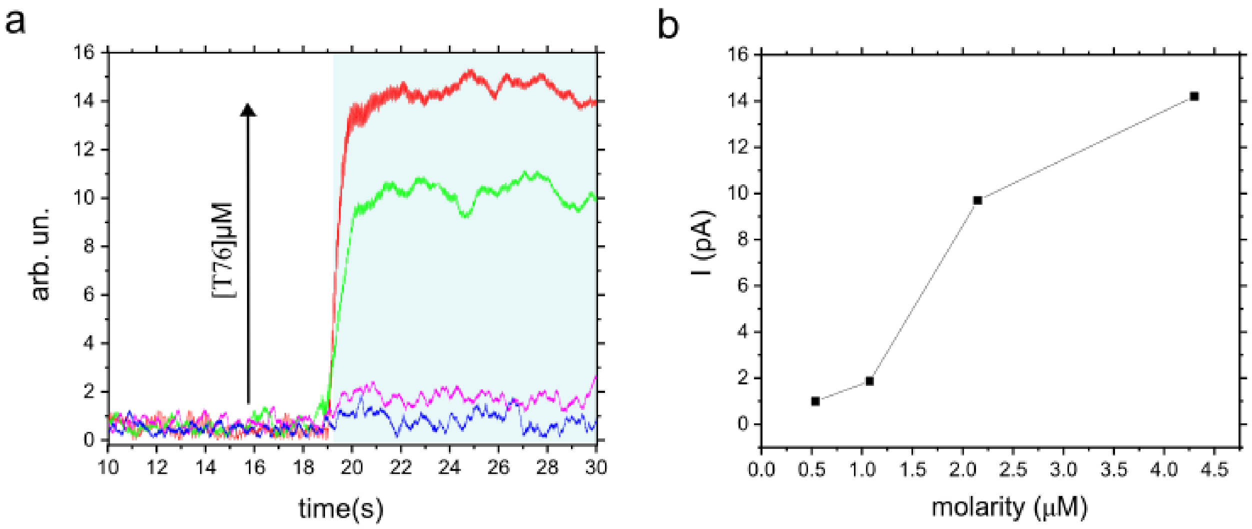


**Supplementary Figure S5** (**a**) Photocurrent traces recorded at increasing concentrations of TARA76-NDs (0.537 μM, 1.075 μM, 2.15 μM, 4.3 μM) under 488 nm laser irradiation (1 mW). The current was measured using the lock-in amplifier (laser modulated at 12 Hz). (**b**) Concentration-dependent photocurrent response corresponding to data shown in panel (a).

***Contribution of Tris/HCl on TARA76 functionality***

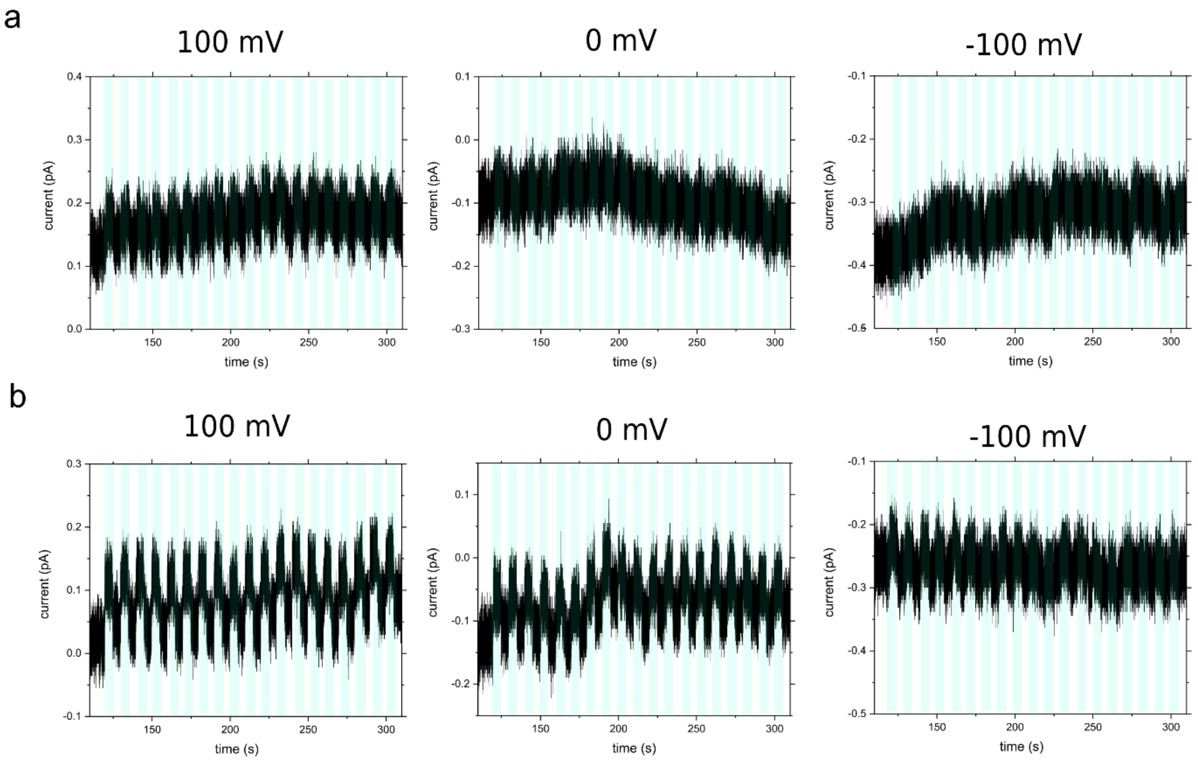


**Supplementary Figure S6** (**a,b**) Current trace recorded after adding TARA76-NDs in the upper chamber of the microfluidic device filled with the unbuffered solution 20 mM Tris/HCl at pH 6.8 (**a**) and 20 mM Tris/HCl at pH 7.6 (**b**), applying a DC voltage of 100 mV (left), 0 mV (middle) and -100 mV (right). Laser light exposure is indicated by blueish time windows.

***Contribution of $Na^+$ on TARA76 functionality.*** Measurements were performed in both buffer and unbuffered solutions at three different applied voltages. Here, we focused on the results obtained with the unbuffered 30 mM NaCl solution (Supplementary Fig. S7a), where we observed that the light-modulated current remains similar at 100 mV and 0 mV. This is consistent with a light-driven proton pump. A comparable trend is observed in the presence of 1 mM HCl added to the 30 mM NaCl solution (Supplementary Fig. S7b), although a distinct initial transient appears consistently in this condition. Finally, in the buffered solution composed of 20 mM $NaH_2PO_4$/ $Na_2HPO_4$ at pH 5.8 with 10 mM NaCl (Supplementary Fig. S7c), a more pronounced peak is observed, along with a significantly higher overall signal. In this case, the transient is consistently present in the same direction regardless of the applied voltage, while the steady-state current varies with negative bias, indicating a voltage/dependent modulation as previously described.[5]

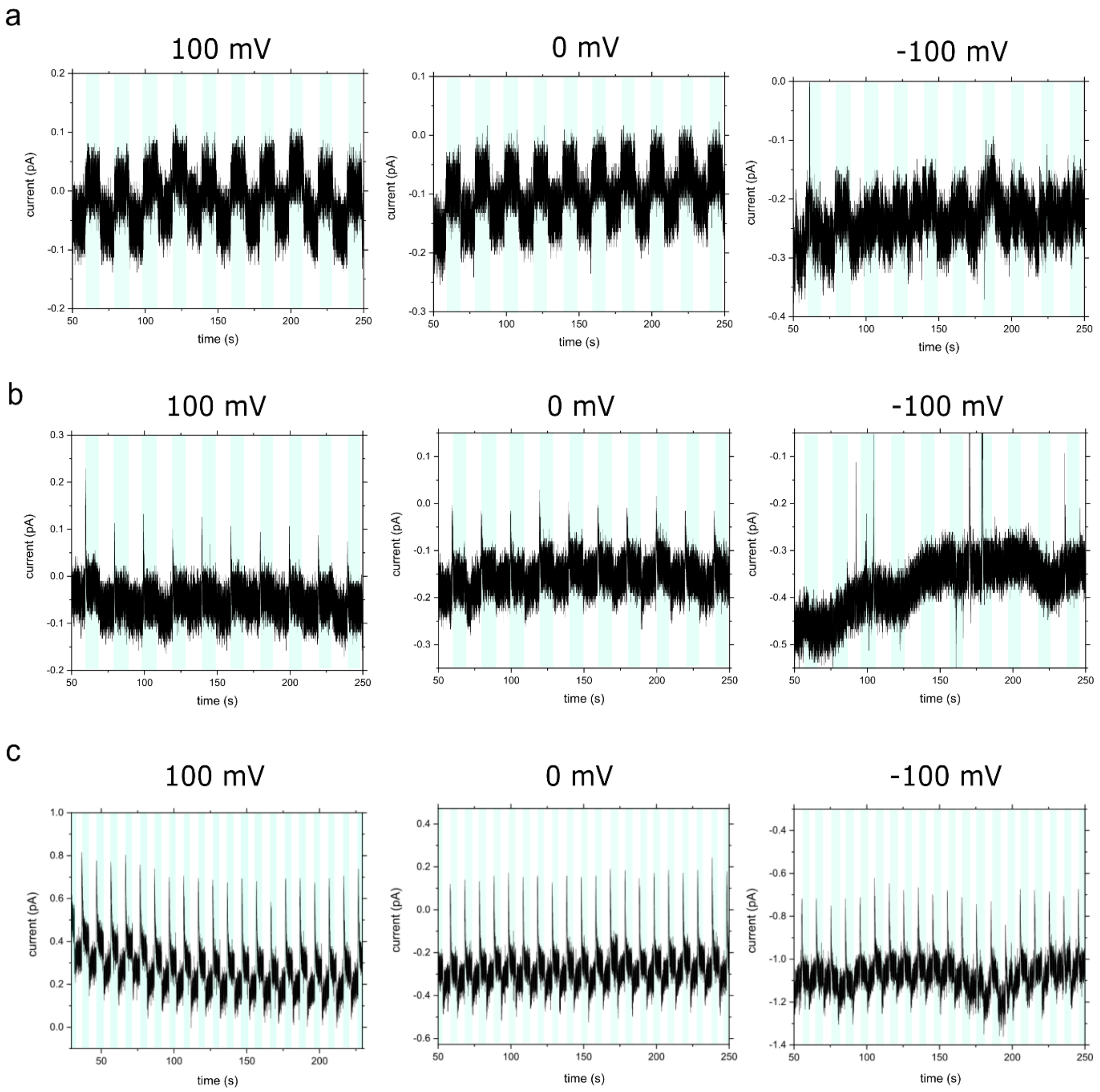


**Supplementary Figure S7** (**a-c**) Current trace recorded after adding TARA76-NDs in the upper chamber of the microfluidic device filled with the unbuffered solution 30 mM NaCl (**a**), 30 mM NaCl + 1 mM HCl (**b**) and 20 mM of $NaH_2PO_4$/ $Na_2HPO_4$ at pH 5.8 + 10 mM NaCl (**c**) applying a continuous voltage of 100 mV (left), 0 mV (middle) and -100 mV (right). Laser light exposure is indicated by blueish time windows.

***Equivalent electric circuit***

The complete model is rather complex, involving resistive element for the solution ($G_{sol}$), and resistive and capacitive elements for the electric double layer ($G_{EDL}$, $C_{EDL}$), the membrane itself (bilayer lipid membrane, $G_{BLM}$, $C_{BLM}$) and finally the contribution of the nanodiscs with TARA76 ($G_{ND}$, $C_{ND}$ in parallel to the membrane). The resistive elements are indicated by their conductance equivalent $G = 1/R$. The order of magnitude of some of these elements have been evaluated by independent measurements with the sole solution, solution and the membrane, and with and without either empty or nanodiscs. Their typical orders of magnitude are reported in the caption to Supplementary Fig. S8, although they are subject to a high degree of variability and to a certain drift over time, as previously stated. Notably, all the additional elements only weakly affect the current response value at the measuring instrument ($G_{meas}$), while they don't affect its shape. The most significant elements are those appearing in the reduced model of the main Fig. 5, namely the conductance $G_{BLM}$ and the capacitance $C_{BLM}$ of the membrane (with or without the nanodiscs), together with the electric double layer conductance $G_{EDL}$. The $G_{ND}$ and $C_{ND}$ sum up to those of the BLM being in parallel, but they cannot be really separated and are shown here only for completeness. In fact, resolving their individual contributes, would require independent measurements with given, known NDs surface occupancies of the membrane but in the very same conditions for all the other parameters. Unfortunately, starting from the membrane surface area itself, there is a certain degree of variability among individual experiments limiting the comparison of total and specific contribution of BLM and NDs incorporated in parallel to each other. Furthermore, we are interested in the effects induced by light exposure, identified in a passive conductance variation and an active current, respectively indicated as $\Delta G$ and $I_{light}$ in the essential (main text Fig. 5) and complete (Supplementary Fig. S8) equivalent circuit. The former is expected to change the slope of the I(V) branches, while the latter is expected to vertically and rigidly shift the whole I(V) curve. The possible variation of capacity $\Delta C$ due to light activation is not represented in the scheme of the circuit because we observed no photo-modulation of such a quantity (apart from a slow drift).

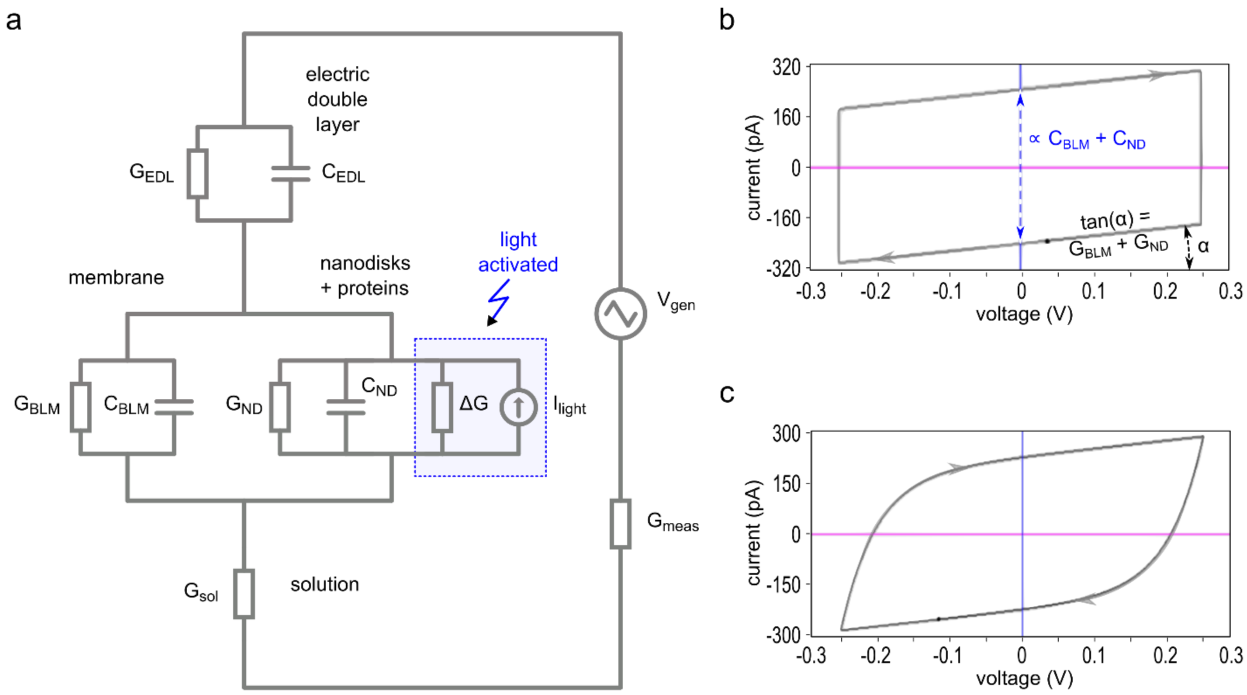


**Supplementary Figure S8** Complete equivalent circuit. (**a**) The circuit has elements associated with the solution, the membrane incorporating empty or protein-loaded nanodiscs, and the electric double layer at its surfaces. The light-induced effects are schematized as parallel conductance variation ΔG and current generator $I_{light}$. (**b**) The parallelogram I(V) cycle is obtained simulating circuit in (a) without the electric double layer resistive element $G_{EDL}$, i.e., letting it be a perfect connection ($R_{EDL}$ = 1/$G_{EDL}$ = 0). (**c**) The same circuit with the presence of a finite resistive $G_{EDL}$ element features an exponential response (rounded edges in the 2nd and 4th quadrant). Simulations in (a,b) were performed by using the online Circuit Simulator tool by P. Falstad [https://www.falstad.com/circuit/]. The used values for the elements in the equivalent electrical circuit are as follows: $G_{EDL}$ = 5 nS, $C_{EDL}$ = 0.1 pF, $G_{BLM}$ = 200 pS, $C_{BLM}$ = 20 pF, $G_{ND}$ = 50 pS, $C_{ND}$ = 5 pF, $G_{sol}$ = 10 µS, ΔG = 20 pS, $I_{light}$ = 0.5 pA.

***AC measurements at different modulation voltages.***

The AC response of TARA76 and its photo-modulation were also evaluated at other triangular voltages, as shown in Supplementary Fig. S9. Here we used seven different $V_m$, starting from 100 mV up to 400 mV, in steps of 50 mV. All the cases do not differ substantially from the results already shown in Fig. 5 for a modulation voltage of 250 mV. As anticipated, such a value represents an optimal choice for $V_m$. Higher modulating voltages instead reduce the noise in the extracted fit parameters over time. However, they also gradually lead to a nonlinear bend in the I(V) cycle, ascribed to dynamical membrane elastic deformation under electrostatic forces between its two sides, or to differential capacity effects. The various sweeps shown in Supplementary Fig. S9 hence allow the optimisation of the modulating voltage. This means being able to select the highest $V_m$ possible (which provides less noise) avoiding significant nonlinear bending of the cycles (with respect to the simplified behaviour).

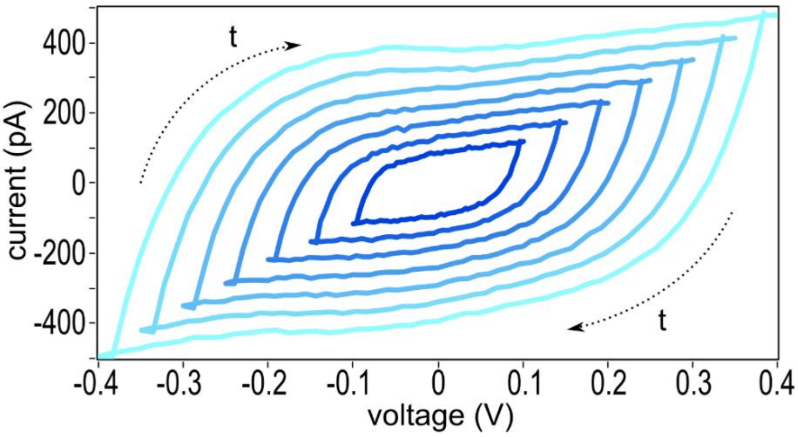


**Supplementary Figure S9** I(V) characteristic obtained by applying different triangular modulating voltages ($V_m$ from 100 mV to 400 mV in steps of 50 mV, corresponding to curves from the inner cycle to the outer one).

In Supplementary Fig. S10 we show the separated conductance ($S^+$ and $S^-$, top and bottom subpanels respectively, in panel a), taken from the fit of the two positive and negative half-wave branches, at three modulating voltages $V_m$ (250 mV, 300 mV and 350 mV). The results are qualitatively the same. On the long-time scale, each conductance appears to be undergoing a slow drift since beginning of monitoring. Their opposite drifts almost compensate each other. At the start of light steps exposure (t = 60 s), they invert such a drift. While the positive conductance shows a slow increase during cyclic light exposures, the negative conductance shows a gradual decrease in time reaching a plateau in approximately 120 s. Both the initial opposite drift and the opposite response to light can indicate an ongoing build-up of an imbalance between the two EDLs in contact on the two sides of the membrane, that is reversed and apparently stabilized by cyclic light exposure. Any effect on the membrane surface area is instead supposed to affect the two conductance in the same proportional way.

We are here interested in the light-induced modulation of conductance on the fast time scale, as shown in the Supplementary Fig. S10c (intermediate time zoom of the previous panel a). The modulation of both S+ and S- are positive in every 5 s time-window of exposure. However, while the steps in $S^+$ are less than 10 pA and their visibility limited by noise and fluctuations, the steps of $S^-$ increase are neatly distinguished and in the order of 25 pA. This is a hallmark of vectorial behaviour in the protein photoresponse and of preferential orientation in its incorporation into membrane. The $\Delta$G amount is almost constant for the different applied $V_m$ and in any case not proportional to the absolute $S^-$ value. A slow drift in time is present also in the capacity extracted from fit (Supplementary Fig. S10b, top). However, while the initial drop/increase of G in the first 60 s is around 10%, the capacity drifts of less than 0.2 pF, i.e. <1% of its absolute value, in the same time range. If this effect is attributable to membrane area, it may indicate a relatively stable surface area.. The fast capacity variations $\Delta$C, i.e. in each exposure window, are instead very weak, at the limit of visibility (Supplementary Fig. S10d, top).

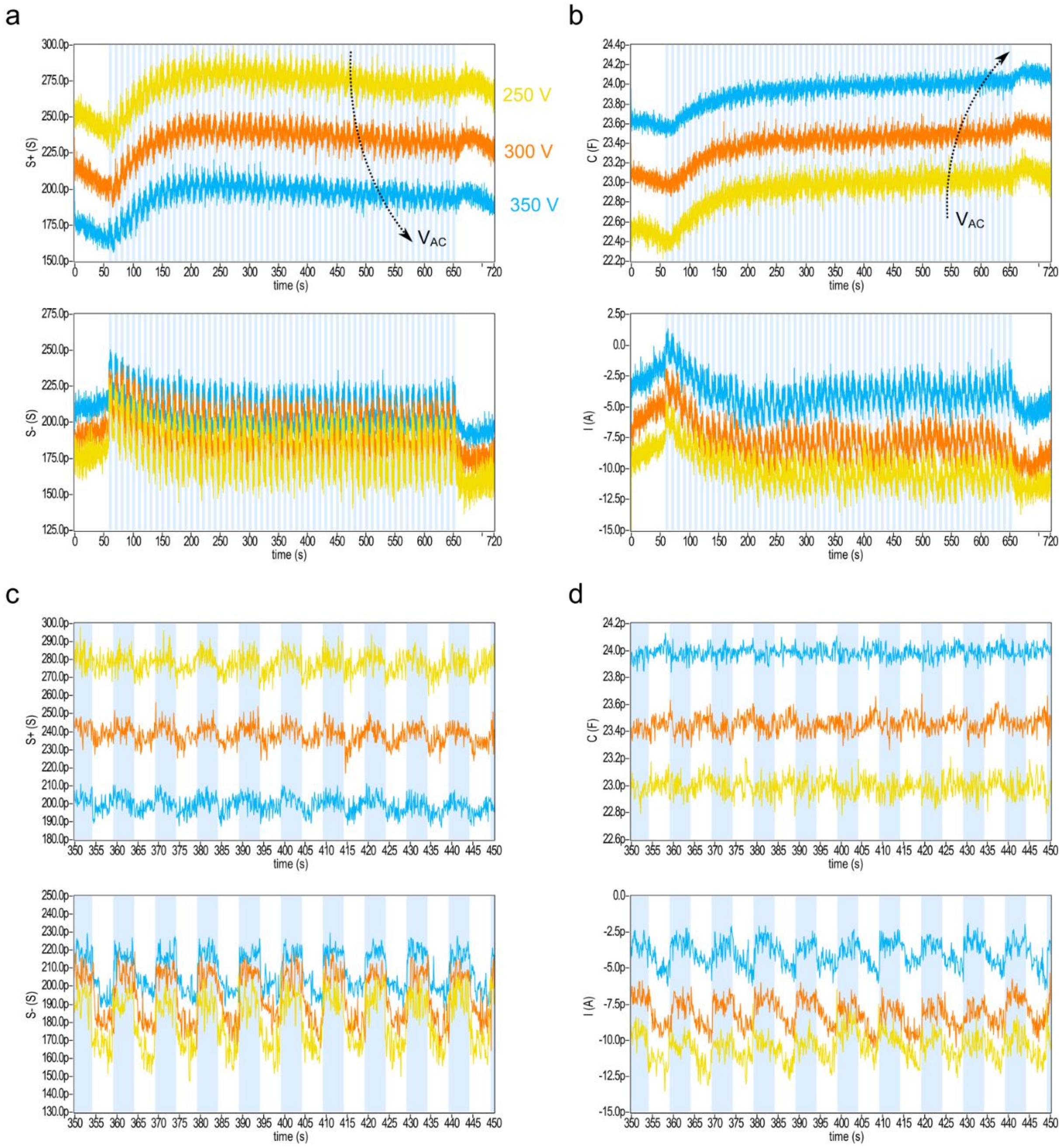


**Supplementary Figure S10** (**a**) Conductance time traces taken from the fit of the positive half-wave (top subpanel) and negative half-wave (bottom subpanel) ramps of the current waveform in presence of TARA76-loaded NDs. The applied triangular voltage at 10 Hz was 250 mV (yellow), 300 mV (orange) and 350 mV (blue curve). The 488 nm laser exposure is indicated by shadowed areas (5 s light on, shaded blue areas, and 5 s light off, white areas). (**b**) Capacitance time traces from the fit of the current waveform (top subpanel). Active current $I_0$ time traces taken from the fit of the current waveform (bottom subpanel). (**c**) Zoom of the light-dependent conductance time traces (100 s time range) taken from the same traces for the panel a. (**d**) Zoom of the light-dependent capacitance (100 s time range) and active current time traces, expanded from panel b. Time step in each panel is 100 ms.

We also studied the zero-voltage (i.e., shunt-circuit) current $I_0$ taken at the crosspoints of the I(V) cycle with the vertical V=0 axis, in each of the two branches. The quantity $I_0 = [I^+(V=0) + I^-(V=0)]/2$ can also be extracted as the average of the two linear intercepts from the fit. Apart from its slow drift (Supplementary Fig. S10b, bottom), a fast modulation of this parameter can be associated with an active current component, reflecting the protein's role as a current generator (i.e., a proton pump). In fact, a fast light-dependent modulation at each exposure was observed (Supplementary Fig. S10d, bottom). Notably, the light-induced changes of the active current are positive. This is paired with the increase in negative conductance (panel c, bottom). Basically, these two quantities appear to balance each other: an active current in one direction and a passive flow in the opposite direction. We think that an important observation stems from this positive modulation of $I_0$ and of negative conductance S-. Basically, the total or average current could appear not being photo-modulated, even in the case of ongoing photo-modulation of the protein and its vectorial activity, The traces in Supplementary Fig. S11 report the active current $I_0$ (dark green trace) and the average current $I_{av}$ during each modulating cycle (light green trace), for the case of $V_m$ = 250 mV. The AC average current does not exhibit evident photo-modulation, apart from some sparse fast peaks. Such peaks are analogue to the peaks seen in the DC transients, but here cannot be resolved due to the time step for one modulating cycle being 100 ms. These results are consistent with the behaviour of a light-driven proton pump, which acts as a current generator by actively converting light energy into ionic flow, nevertheless subject to a passive ionic counterflow. The balance of these two flows depends on the specific conditions, the very same as the stationary current in DC can even get to zero after a transient, depending on environmental conditions. Hece, an ongoing vectorial photoactivity of the protein cannot be excluded only by observing a null stationary DC or average AC current. The combination of the response in DC and AC conditions may provide valuable insights into both the protein's extended orientation and its functional role as a light-activated proton pump, which not only generates an oriented active current but also shows a directional photo-conductance upon illumination.[6]

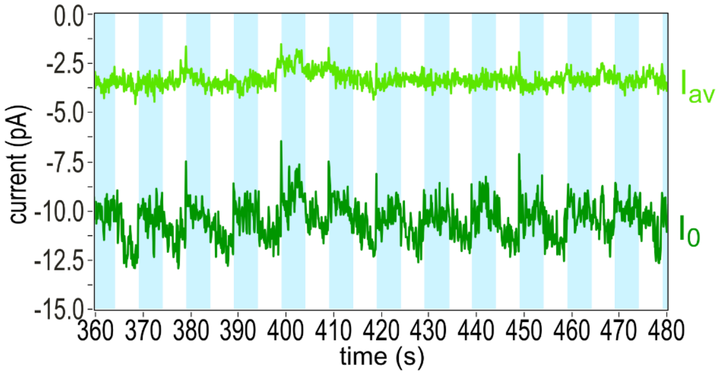


**Supplementary Figure S11** Time traces of $I_0$ and $I_{av}$ from the I(V) cycle at $V_m$ = 250 mV. The active current is defined as $I_0 = [I^+(V=0) + I^-(V=0)]/2$, while $I_{av}$ represents the average current in each 100 ms cycle at 10 Hz.

***TARA76 orientation into the artificial lipid membrane.*** If the membrane is stable for a long time, it is possible to register the photocurrent at different measuring DC voltages in the same experimental sequence. As it is shown in Supplementary Fig. S12a,c and e, the photocurrent peak has same direction and is independent on the measuring voltage, respectively at a DC voltage of +80 mV, 0 V and -80 mV, confirming the presence of a proton-pumping activity that is oriented by the initial voltage of +80 mV, also applied during proteins-loaded NDs incorporation. The magnifications over 4 on/off steps for each measurement are shown on the right part of the respective panels. On the contrary, the stationary current, is in accordance with the orientation of the DC voltage that is applied during measurements (Supplementary Fig. S12b, d and f), despite very small with respect to the peak.

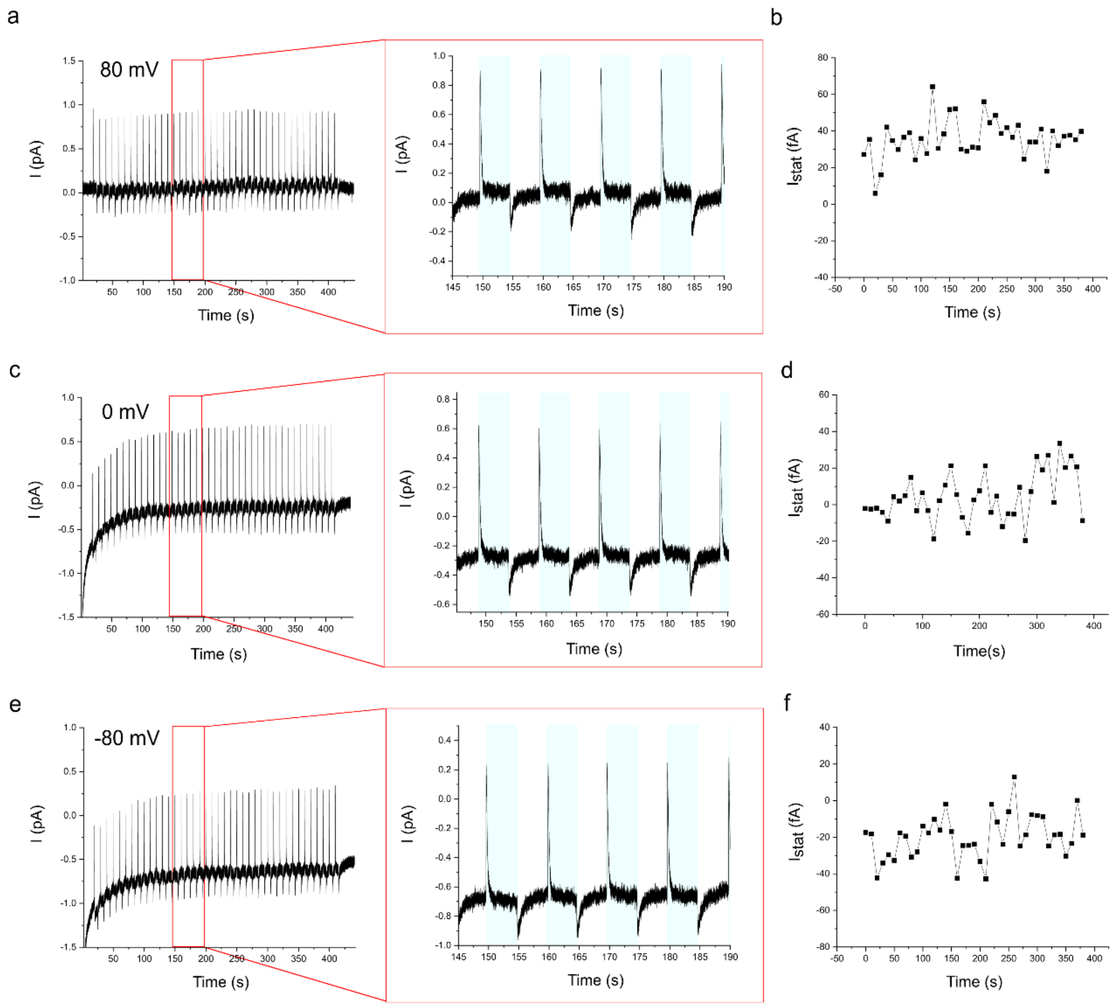

**Supplementary Figure S12** (**a**) Current trace recorded at continuous voltage of +80 mV after the addition of TARA76-NDs to the upper chamber of the flow cell (right subpanel shows the trace magnification in a range of 45 s). (**b**) Stationary current $I_{st}(t)$ during time extracted considering the TARA76 light-induced plateau current, for each exposure window in panel a. (**c**) Current trace recorded at a continuous voltage of 0 mV, after the sequence in panel a. (**d**) Stationary current $I_{st}(t)$ during time extracted from the trace in c. (**e**) Current trace recorded at a continuous voltage of -80 mV after the sequence in panel b. (**f**) Stationary current $I_{st}(t)$ extracted from the trace in e. All acquisitions were performed without any membrane breaking between the different sequences.

Differently from the cases above, we now applied different DC voltages during the membrane formation in presence of TARA76-loaded nanodiscs. In the first case, TARA76-NDs were added on the flow cell before the artificial membrane formation, and we kept applied a continuous voltage of +80 mV. After 20 min of incubation the artificial membrane with TARA76-NDs was irradiated with on/off cycles of 5 s, using a 488 nm laser at 4 mW. As we can see from the Supplementary Fig. S13a the initial peak in all the transients is positive, indicating an orientation of the protein activity, co-directional with the voltage applied during incorporation and measurement. The stationary current has also the same direction being around +0.1 pA (panel b). After this measurement the membrane broke up and was formed again, but now keeping applied an opposite voltage of -80 mV. After the formation and the incubation for 20 min, we started cyclic irradiation (5 s light on). We noticed a change in the polarity of the photocurrent, in which both the initial spike transient Supplementary Fig. S13c and the stationary current of -80 pA (panel d) are oriented in the opposite direction than before. Subsequently, the membrane was broken up again and was reformed keeping applied a +80 mV voltage. After 20 min of incubation, cyclic light-exposure started again as shown in Supplementary Fig. S13e. The signal reverted to its initial polarity, giving positive orientation of both the transient peak and the stationary current at +140 pA (panel f). Combining the results from the Supplementary Fig. S12 and Supplementary Fig. S13 we can conclude that the DC voltage applied during membrane formation with simultaneous presence of TARA76-NDs, can determine the direction of the transient peak at the switch-on of light, while the voltage applied during measurements affect the stationary current. Hence, application of a DC voltage during membrane formation represents a valid strategy as forcing mechanism able to aid incorporation of TARA76-NDs with a strongly preferential orientation.

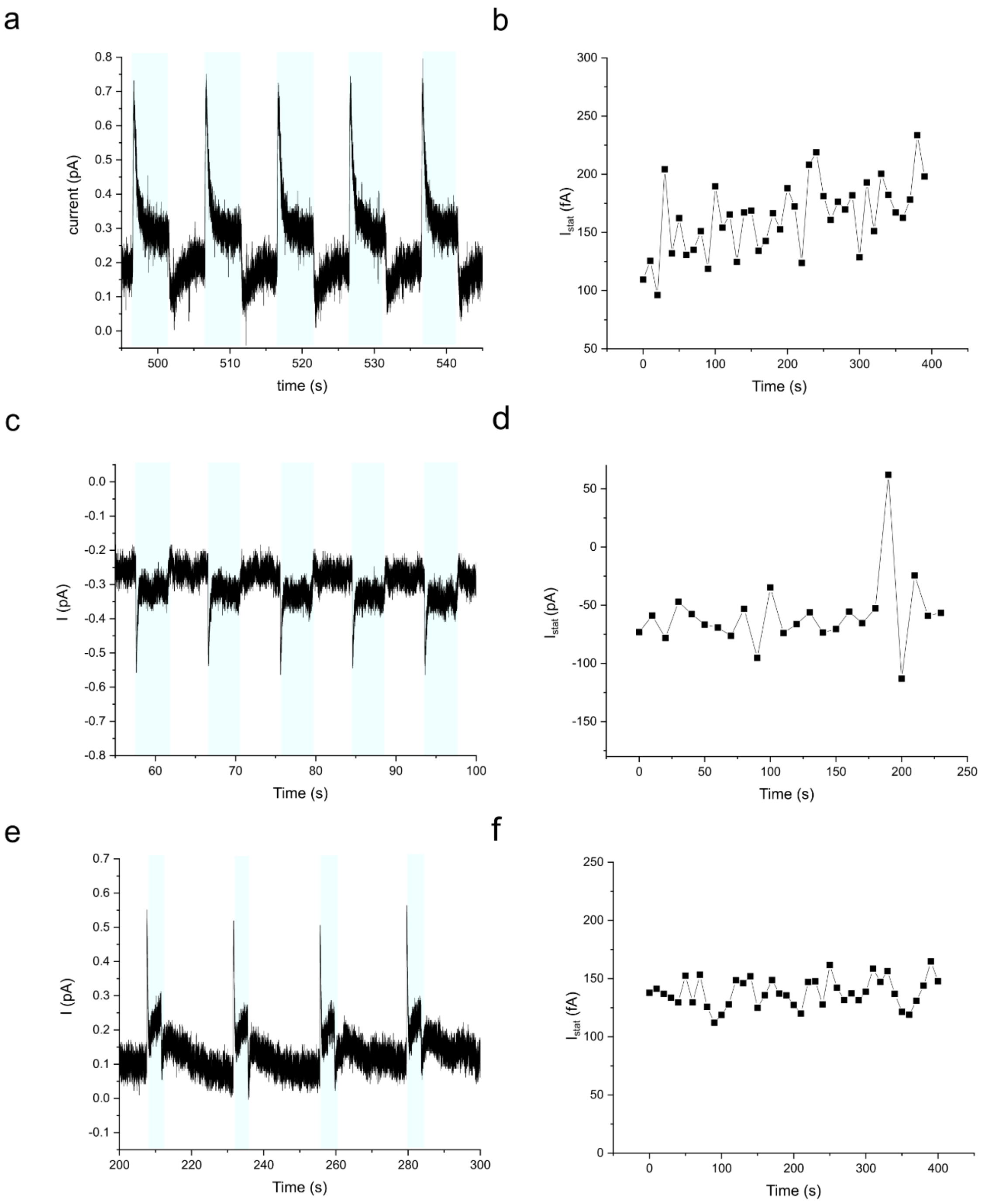


**Supplementary Figure S13** (**a**) Representative current trace recorded at continuous voltage of +80 mV after the addition of TARA76-NDs to the upper chamber of the flow cell. The artificial lipid membrane was formed in the presence of the protein. (**b**) Curve extracted considering the TARA76 light-induced stationary current at each exposure window of the trace shown in panel a. (**c**) Current trace recorded at a continuous voltage of -80 mV in presence of TARA76-NDs during membrane formation. (**d**) Curve extracted considering the TARA76 light-induced stationary currents corresponding to the trace shown in panel c**.** (**e**) Control current trace recorded at continuous voltage of +80 mV in the presence of TARA76-NDs during artificial membrane formation. (**f**) Curve extracted considering the TARA76 light-induced stationary current corresponding to the current trace shown in panel e.
